# Reciprocal Environmental Decision Support (REDS): better tailored advice in return for data

**DOI:** 10.64898/2026.01.08.697712

**Authors:** Ben Kenward, Nick Casey, Perline Bastid, Francis Buner, Ishi Buffam, Julie Ewald, Robert Kenward

## Abstract

1. Environmental Decision Support systems provide model-based predictions, tailored to user-inputted information about local ecosystems, which can support management decisions by citizens. Citizen Science systems accept user input to improve models. Thus, each system type emphasises automatic data transfer in one direction: model to citizen or citizen to model, respectively.

2. We introduce a new system type that combines automated data transfer in both directions.

3. Reciprocal Environmental Decision Support (REDS) systems process user-contributed information to (1) provide tailored predictions supporting management, and (2) improve the underlying predictive model.

4. For our proof-of-concept REDS system, Garden Advice, we began with a Bayesian species-habitat association model (for the House Sparrow *Passer domesticus*) based on existing data, and obtained new data from UK residents about habitat structure and sparrow observations in domestic gardens. The model made predictions about sparrow presence, and effects of planned garden changes. Model parameters were updated by contributed data; parameter estimates generally tightened. One notable update likely reflects reality (a positive association with grass), while another (a reduced association with roof proximity) likely reflects observation bias.

5. The updated model predicted the new data better than the original model. Thus, untrained observers can provide data of sufficient quality to refine a model of trained observer data. Notwithstanding important questions about distinguishing observation bias from ecologically meaningful information, our system demonstrates that important synergies can be obtained from the REDS approach. Later users of the system obtained better advice thanks to automatically incorporated contributions from earlier users.

**Lay Summary:** A summary for non-specialists is available at https://bit.ly/reds_brochure.

## Introduction

### Introducing Reciprocal Environmental Decision Support (REDS)

Environmental Decision Support (EDS) systems support environmental management by providing model-generated predictions about environmentally relevant consequences of possible actions (McIntosh et al., 2011; Wong-Parodi et al., 2020).^1^ The value of EDS systems lies in their ability to provide advice tailored to the specific circumstances of the user, modelling the impact of potential management changes. EDS systems therefore typically require complex query input, because optimised predictions about environmental consequences of decisions require the ecosystem and its context to be described in some detail.

Frequently, land managers collate and input detailed information, that is processed to produce decision-support output, but is not put to further use. As one example illustrating this general point, the Cool Farm Tool uses farm activity data to estimate farm carbon footprint and to support decisions by analysing hypothetical activity changes; it has been used for over 125,000 assessments (Cool Farm, 2025a; Kayatz et al., 2019). This implies a rich central database of data relevant for ecosystems on farms, but as they note, “the Cool Farm Alliance has not used any data stored in the tool” for purposes except returning assessment results (Cool Farm, 2025b). This stands in contrast to many other information systems, where value is obtained from query input (e.g., Bondia-Barceló et al., 2016), with useful examples including web search data revealing trends in public consciousness of environmental issues (Nghiem et al., 2016) and enabling epidemics to be rapidly tracked (Ginsberg et al., 2009).

The richness of EDS query input data, along with the lack of evidence for its use beyond returning query output (we explore relevant evidence below), imply that important opportunities are being missed. The typically multivariate nature of this data implies that it could parameterise the predictive models used within EDS. This would represent the incorporation into EDS of Citizen Science (CS), in which non-expert citizens contribute data that updates models which can be used to generate advice (Fraisl et al., 2022; Johnston et al., 2025; McKinley et al., 2017).

The current work introduces a new type of system that automates the combination of EDS and CS in this way (Kenward et al., 2013; Papathanasiou & Kenward, 2014). Whereas CS systems often automate the one-way data transfer from citizen to system, and EDS systems automate the one-way data transfer from system to citizen, Reciprocal Environmental Decision Support (REDS) represents a novel system type which automatically closes this loop (Figure 1). REDS systems are EDS systems that are reciprocal; not only is user knowledge updated by the model predictions, but model predictions are automatically updated by user observations (Figure 1).

**Figure 1.**
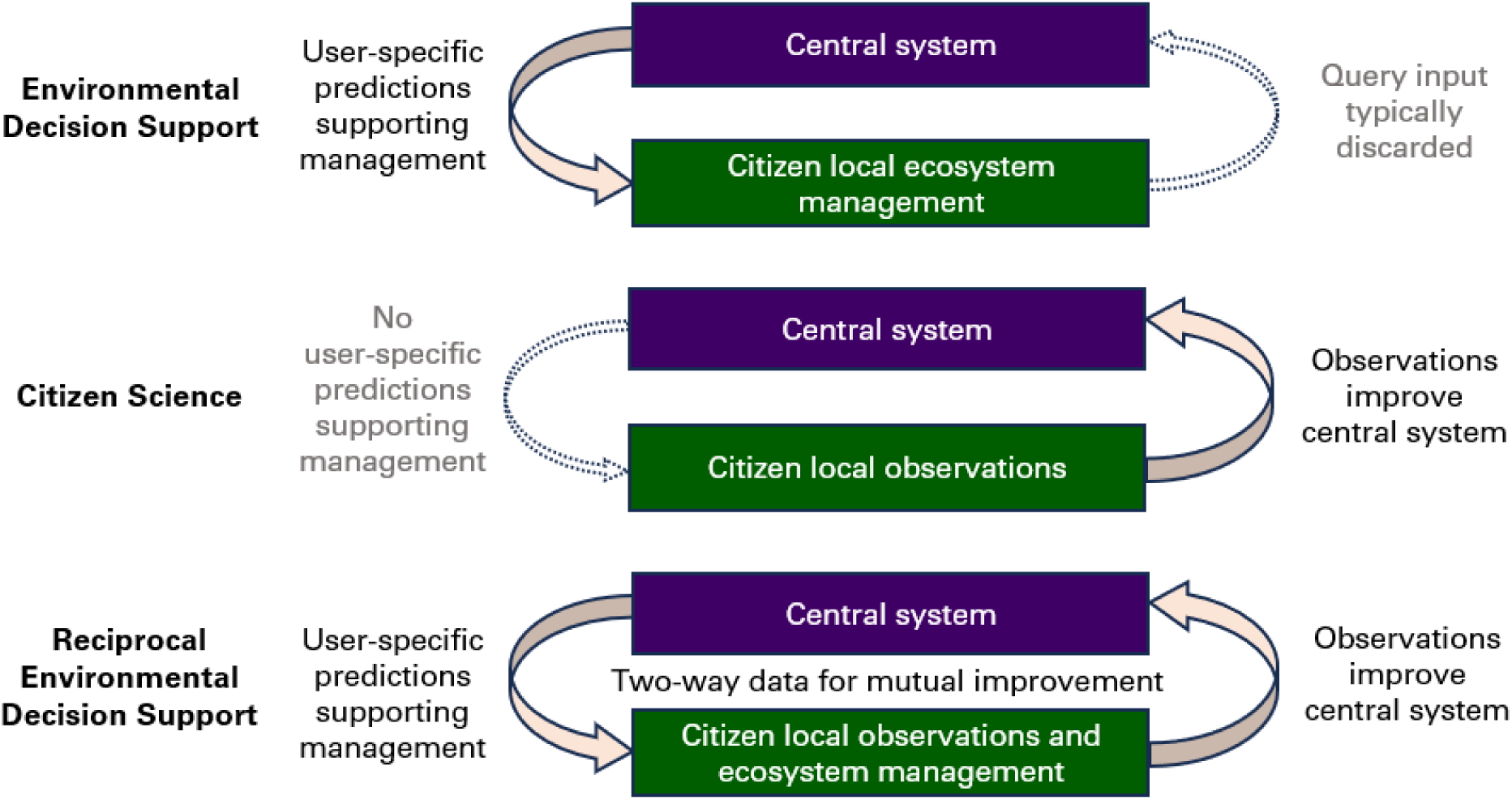
Automated information flows in EDS, CS, and REDS.

Automated information loop closure has the theoretical potential to revolutionise the quality of data, models, and decisions, through iterative mutual improvement via feedback loops (Wiener, 1948). The automated closure of the EDS / CS loop is important because it allows new information to quickly influence tailored advice to specific users. This is analogous to the way that a travel-planning app can use real-time information from other users to help travellers avoid delays. Currently, EDS models may be parameterised with data from CS, but this takes months or years; reports from CS programs can influence management behaviour, but on a similar time scale, and without being easily adapted to individual users’ specific circumstances.

Similar arguments have prompted the application of the digital twin (DT) concept to environmental science. DTs are “digital counterparts of a physical object or process” that is “continuously updated with data, thereby representing the state of something in current time” (de Koning et al., 2023). DTs close information loops to rapidly update models, but do not necessarily acquire data from system users, and do not necessarily provide EDS. We summarise key differences between the definitions of CS, EDS, REDS, and DT systems in Table 1.

**Table 1.**
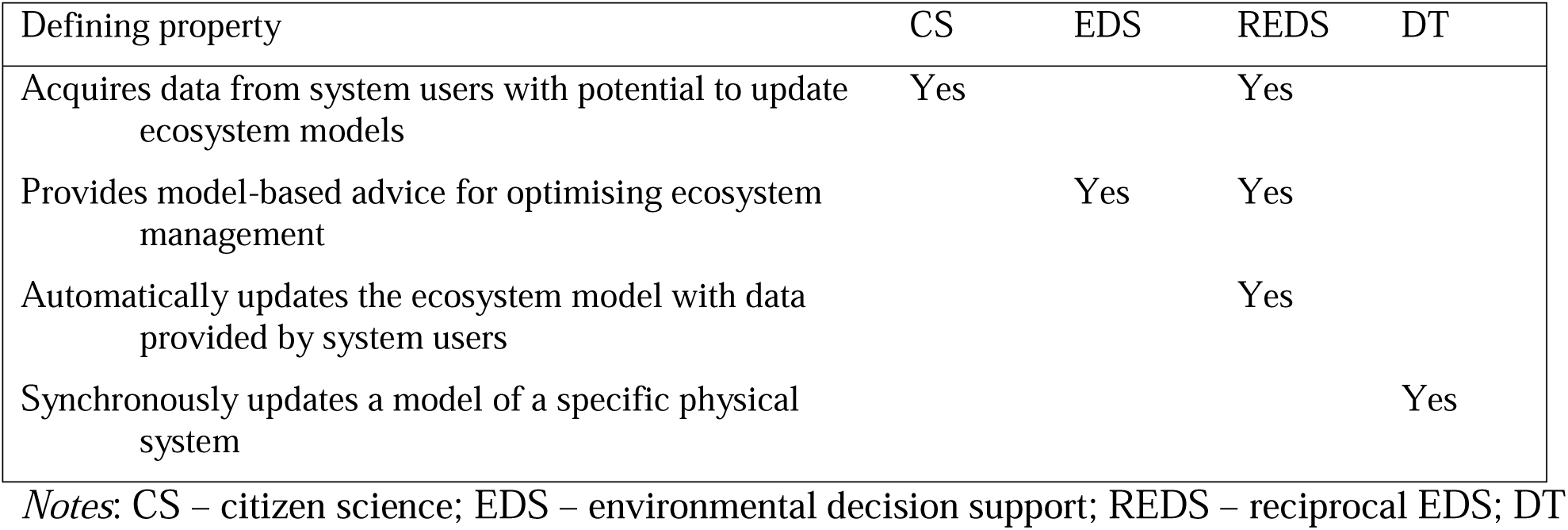
Defining properties of environmental data systems (Yes = required by definition)

Here, we empirically demonstrate a REDS system. It uses input not only to generate model predictions that can support decision making, but also to improve the model itself, on a time scale of days. We illustrate this by implementing a habitat association model for an animal indicator-species that makes predictions about species occurrence based on user-provided habitat descriptions, with model updating enabled by users also providing animal observations.

### Searching for existing REDS systems

Some systems feature reciprocal sharing of environmentally relevant data but are not REDS (or EDS) because there is no model, for example the PurpleAir system for sharing air quality measures (PurpleAir, 2025; Robinson et al., 2023). The species-mapping functionality of iNaturalist does have a model that is reciprocally updated: users report species observations that update a model predicting occurrence at unobserved locations (Cole et al., 2023). The data flow and model updating is REDS-like, but iNaturalist is not an EDS. The species occurrence models are based on spatial coordinates at very coarse scale, and (as implemented) do not involve environmental conditions predictors, so the models are not suitable to support the what-if analyses needed for EDS. The situation is similar for other popular CS platforms such as eBird. Although eBird does operate on a spatial resolution that can be useful for management decisions (Farwell et al., 2026), it does not offer users predictions about the consequences of modelled changes to local land management.

Google Maps route-planning is presented as supporting environmental decisions, offering information on the relative energy efficiency of different routes. Further, it is reciprocal, as measurements of traffic speed from users update its models of traffic conditions which in turn inform route recommendations. Thus, Google Maps route-planning is the most REDS-adjacent system of which we are aware (although perhaps not a successful one, Burris et al., 2023). It stretches the definition, as it is reciprocal and claims environmental benefits, but does not assist ecosystem management directly.

Despite recent reviews of DT systems in environmental science (Hazeleger et al., 2024; Trantas et al., 2023), we are unaware of any DT system yet operating practically to provide decision support for ecosystem managers based on predicted consequences of potential management choices. Many systems aiming to do so are reported at a prototype stage (e.g., De Antoni et al., 2026; Lu et al., 2026; Taubert et al., 2024). We know of two fully operational systems that are framed as DTs that can automatically exploit user observations – the crane radar, which informs birdwatchers as to where to find migrating common cranes *Grus grus* (De Koning, 2025), and a Finnish app (*Muuttolintujen Kevät*) for detecting and recording birds (Ovaskainen et al., 2026) – but these are not EDS. They do not facilitate users’ ecosystem management by providing them with predictions about the consequences of potential management decisions.

Following a systematic review (see Supporting Information), we found no existing EDS systems with reciprocal updating. We therefore introduce Garden Advice as the first REDS system for ecosystem management.

### The Garden Advice REDS system and our associated research questions

Garden Advice (GA) records (amongst other things) habitat descriptions and House Sparrow *Passer domesticus* observations in domestic gardens across the UK. We chose a sparrow habitat association model for testing REDS-related research questions for multiple reasons. House Sparrows are in decline in many global regions, for complex reasons, and are frequently studied, so are a useful indicator species (Hanson et al., 2020). Despite their decline they are relatively common in UK domestic gardens, thanks to their association with humans (De Laet & Summers-Smith, 2007). UK domestic gardens are an excellent arena for citizen science, thanks to the popularity of gardening (RHS, 2025; Sandhaus et al., 2019; Sharma et al., 2019). Importantly, a raw dataset is available for sparrow habitat association in UK domestic gardens, thanks to the Glasgow House Sparrow Project (GHSP; Matthiopoulos et al., 2019), enabling an initial model.

Our primary research question was whether we could successfully implement a functioning REDS system, defined as confirmation of the following preregistered hypothesis: after updating the initial model with user contributed data, the model will better predict the user data. Our focus is on predicting newly user-contributed data rather than the initial data, because this directly addresses a key prerequisite for a REDS system – the addition of user data must make the system better at predicting user data. In principle, this is a modest criterion, because if there is information in the user data absent from the initial model, then with enough data, logically the criterion must be met. In practice, when soliciting internet data from untrained users across an appreciable geographic range, there are reasons why these conditions may not apply, rendering the question non-trivial. A secondary purpose of this study is to describe user advice-seeking behaviours within Garden Advice, to investigate how those managing land might use a REDS system.

Our methods include no independent verification of accuracy of citizen contributions. This raises concerns about citizen science data quality (Aceves-Bueno et al., 2017; Backstrom et al., 2024; Johnston et al., 2023; Kosmala et al., 2016), questionable in similar contexts without additional verification (Williams et al., 2015). We return to this issue in the light of our results.

## Methods

### Declarations

Extensive additional detail concerning all methods is provided in Supporting Information. Large Language Models were sometimes used to check natural language style and to assist with writing of code. The study received ethical approval from the [ANONYMISED] human-participant research-ethics internal assessment procedure. Participants gave fully informed consent. We preregistered a specific method to test the preregistered hypothesis (Parker et al., 2019), which we followed except where unforeseen circumstances intervened, as noted where relevant.

### Participants

Prolific, a service for social scientists to recruit paid participants, known for good quality data in comparison to other similar services (Albert & Smilek, 2023; Peer et al., 2022), was used to commission 71 UK residents to describe their gardens using Garden Advice in May 2025. An additional 68 UK residents made initial attempts to participate, but could not, for example due to misidentifying birds in a screening questionnaire. All garden data obtained during the study period that could be included in model updating was included. A small number of broad requirements for participation (e.g., having an upper secondary education) were implemented in Prolific to optimise participant ability and motivation to succeed at mapping a domestic garden. Most participants were middle-aged; most were in full-time work; most had university degrees.

### Garden Advice app

The system provides user-friendly methods for citizens to map habitats in domestic gardens and to specify areas where they typically observe House Sparrows. The system is REDS because it provides users with scores indicating the likelihood of observing sparrows in gardens they describe, and updates the model that generates these scores with the user-provided data. The system also indicates how planned garden changes would impact the sparrow scores. Thus, the system is designed to be capable of providing better EDS, the more it is used – the hallmark of the REDS approach. Garden Advice can be accessed at https://garden.earthadvice.org (those providing non-genuine data for testing purposes are asked to enable testing mode).

In-app instructions guided the users to (1) find their garden on the base map, (2) draw a boundary around it, (3) add habitat patches until the bounded area was filled, (4) certify that the map was accurate (with direction to editing functions if not), (5) answer whether they had ever seen a House Sparrow in their garden, (6) if they answered yes, draw polygons around all specific areas where sparrows have been seen. All data was contributed without any non-automated assistance to users.

Garden Advice also incorporates a carbon model estimating how much carbon gardens sequester. It provides these estimates in the form of hours of human life saved, by converting carbon to human mortality via the thousand-ton rule (see Pearce & Parncutt, 2023 for a presentation of this rule, and Bressler, 2021 for an example model from which it is derived). Although not part of the REDS functionality, this is an important part of Garden Advice because it attracts users (this was how we developed the system), and because framing carbon sources and sinks in terms of countable human deaths and lives saved is a useful messaging tool (Kenward, 2024).

### The sparrow habitat association model

#### Variables

Model variables were chosen for compatibility with the publicly available Glasgow House Sparrow Project dataset (GHSP), consisting of observations from trained surveyors at 32 individual colony areas in and around the city of Glasgow (Matthiopoulos et al., 2019). Each data point represents a 2 m^2^ map grid-cell where a sparrow was (1) or was not (0) observed. Predictor variables are proportion of cell covered by each of six habitat types (tree, bush, hedge, grass, roof, and artificial surface) plus two distance variables representing the distance from the cell to the nearest hedge or roof. Garden Advice (GA) allows mapping of two additional habitat types anticipated to be necessary for participants to fully map their gardens: water (ignored in the current model) and cultivated beds (treated as bushes).

The GA app solicits information about sparrow observation frequency in each polygon where sparrows are reported as observed, but this information must be simplified for compatibility with the GHSP data structure. Thus, the GA binary dependent variable is 1 if any sparrow observation area overlaps with the cell, otherwise 0, irrespective of observation frequency in that area. Utilised GHSP distances were approximate reconstructions because available GHSP distances were *z*-transformed (normalised to *M* = 0, *SD* = 1) before release; GA roof distances are calculated using existing map data as well as user data; GA hedge distances in gardens with no hedges are imputed as the maximum observed value (96.7 m). For presentation, distance variables are converted to proximity variables by reversing the sign, so that effects have a more intuitive direction.

#### Model specification and updating

The model is a Bayesian binary logistic regression, with R syntax:

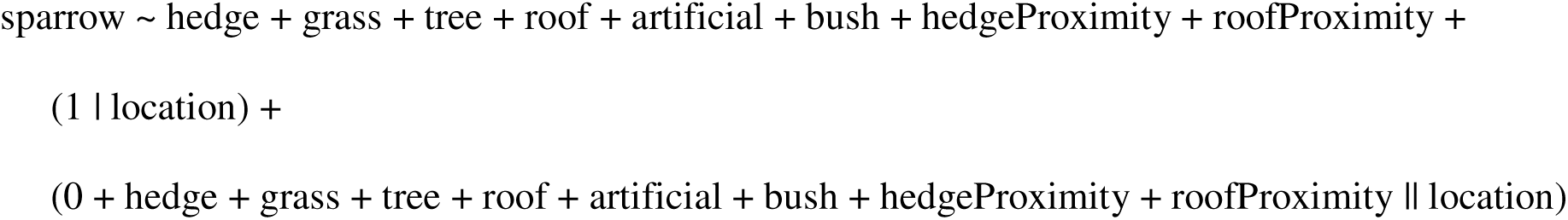

In other words, there are eight fixed predictors, an overall intercept, a random intercept for each observed location (GHSP colony or GA garden), and random slopes for each fixed predictor and location (uncorrelated to ease model fit). The random slopes control for intra-location spatial autocorrelation, as well as capturing any inter-location heterogeneity in habitat associations. We chose a Bayesian approach in part because the REDS approach is inherently Bayesian, as it involves the update of model parameters based on new data.

The model was initially fit to all GHSP data and incorporated into the GA system. It was then updated with the new GA data every day during the study period, entirely automatically except for manual addition of certain data contributions to an exclusion list (Table S3).

GHSP data was sampled in case-control format – each colony area was observed until a sparrow was registered at 40 grid cells, then 40 grid cells where no sparrow was observed were randomly selected, and habitat was described for the 80 cells (Matthiopoulos et al., 2019). GA data, on the other hand, comprises full presence-absence information, enabling prevalence calculations. We used a prevalence-based offset to make the data sets compatible. Information about offsetting in the Supporting Information is important because it involves a deviation from preregistration, for the sake of improved compatibility when comparing models.

Preregistration stated that models for leave-one-group-out cross-validation (see below) would be fitted using the standard Markov chain Monte Carlo (MCMC) approach, but in practice these models ran too slowly so we switched to Integrated nested Laplace approximation (INLA).

Priors were the default in all cases (as preregistered), except for random slope variance hyperparameters in the INLA models, where initial poor fit, combined with re-consideration of appropriate prior beliefs, led to a specification of a comparatively high variance prior.

#### Model predictions and their evaluation

Random components of the model explain large amounts of variation. However, this variation cannot be used for predictions at novel locations, and we therefore only use fixed model components for making predictions. For evaluating prediction accuracy, we preregistered the use of Brier scores (equivalent to mean squared error; lower scores are better) and the standard decomposition into two primary components (Merkle & Hartman, 2018; Siegert, 2017).

Discrimination (also known as resolution) measures how well separated the forecast probabilities are for observed presences versus observed absences (higher scores are better). Miscalibration (also known as calibration) measures how far the forecast probabilities deviate from the observed frequencies on average (lower scores are better). We ignore the third decomposition, uncertainty, which measures uncertainty inherent to the data.

Our primary model comparison examines the relative abilities of the initial GHSP-data-only model, and the model updated with GA data, to predict GA data. This is because the GA data is assumed closer to the population we aim to represent with this model – but we also explore other predictions and discuss this assumption. When comparing model performance metrics, we generate them using leave-one-group-out analysis (Vehtari, 2025), a hierarchical leave-one-out cross-validation where a model is never evaluated for predicting data it is trained with. Our preregistered approach was to leave out one location (garden); as an additional robustness check we leave out one geographical region.

## Results

### Data description

Gardens were distributed throughout the United Kingdom (Figure 2). Descriptive statistics for model variables are in Table S4. Key figures include that sparrows were observed in 85% of GA gardens and in mean 22% of GA map grid cells. Examples of user-created maps are in Figure 3.

**Figure 2.**
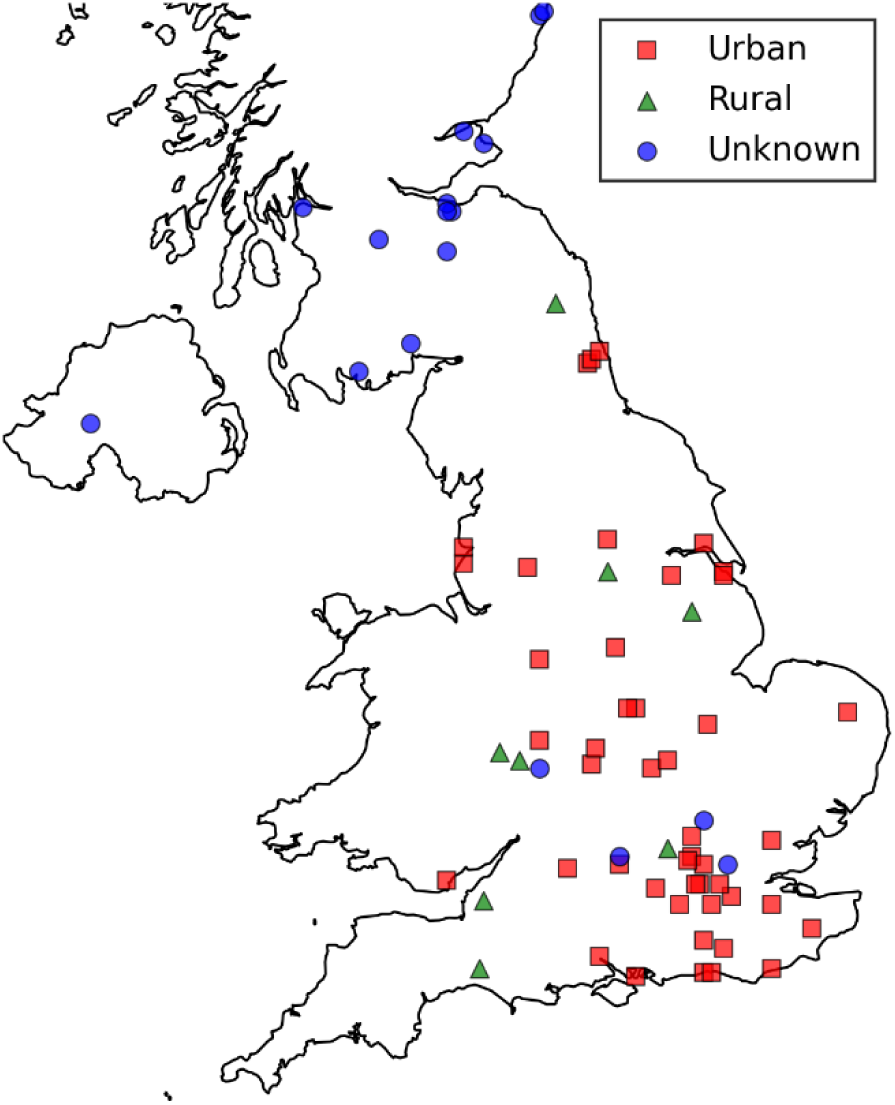
Locations of the 71 Garden Advice (GA) gardens within the UK, snapped to a 5 km grid for privacy. Rural / urban classification data (ONS Geography, 2025) covers England and Wales only and is sometimes unavailable for other reasons.

**Figure 3.**
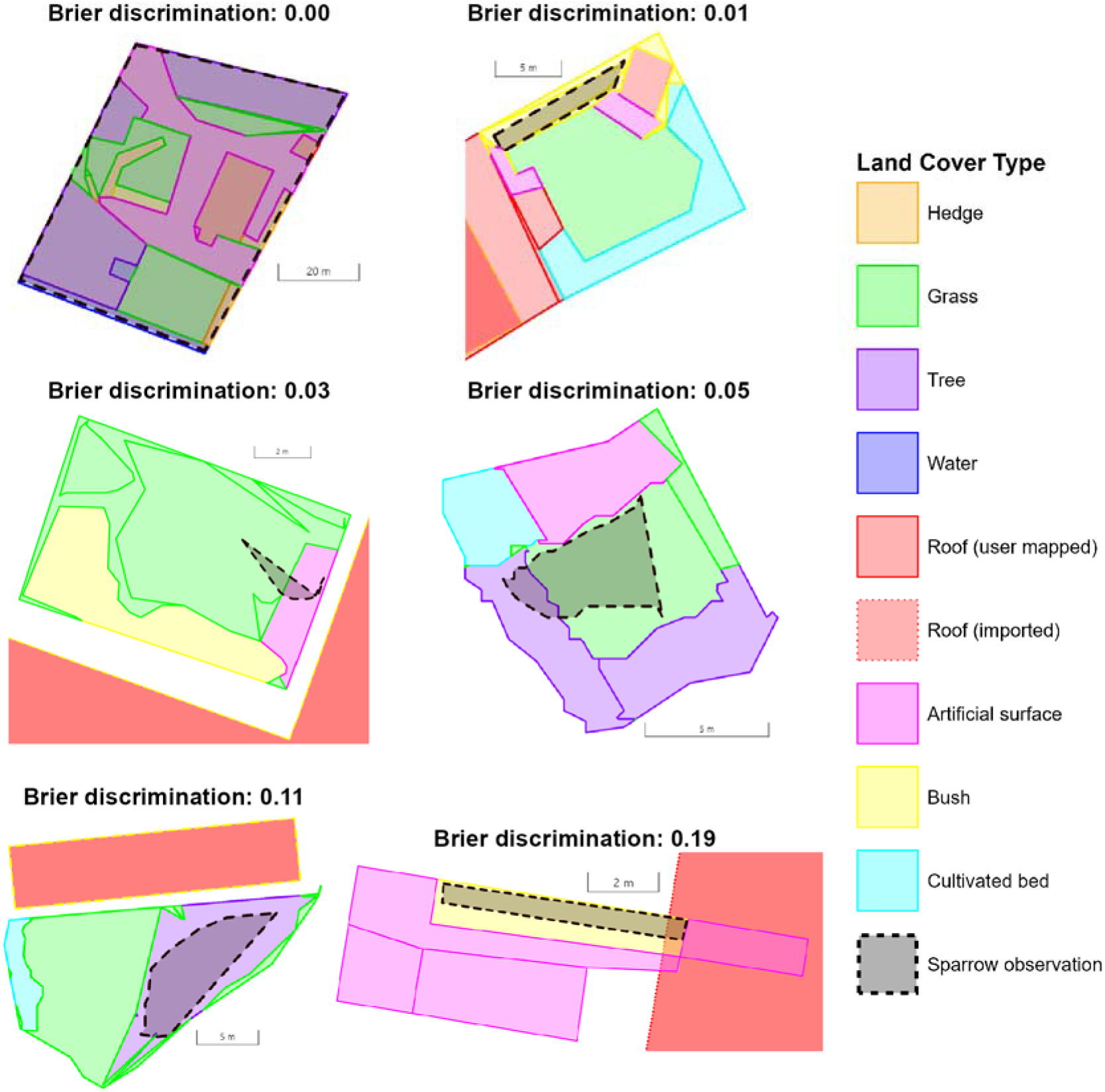
Six user-provided GA garden maps, selected as those nearest the duodeciles (0%, 20%, … 100%) of the Brier score discrimination decomposition distribution, to provide examples of where sparrows are poorly and well predicted by the final model. Buildings with dotted edges are not drawn by users but automatically imported to calculate roof proximity.

### Effect of Bayesian updating with Garden Advice data on model parameters

Figure 4 shows how fixed model parameters were updated through the addition of GA data (and incidentally, indicates very good convergence between INLA and MCMC models – see also Supporting Information Section 4.3.2.1). For trees, bushes, and hedges (cover and proximity), the updated model became more confident of more positive effects. The effect size differences were small, but the model updated from being only weakly confident to being very confident in positive effects. There was a salient change for grass, where the model updated from being weakly confident in a negative effect to slightly more confident in a positive effect (with a 94% probability this represents a real difference^2^).

**Figure 4.**
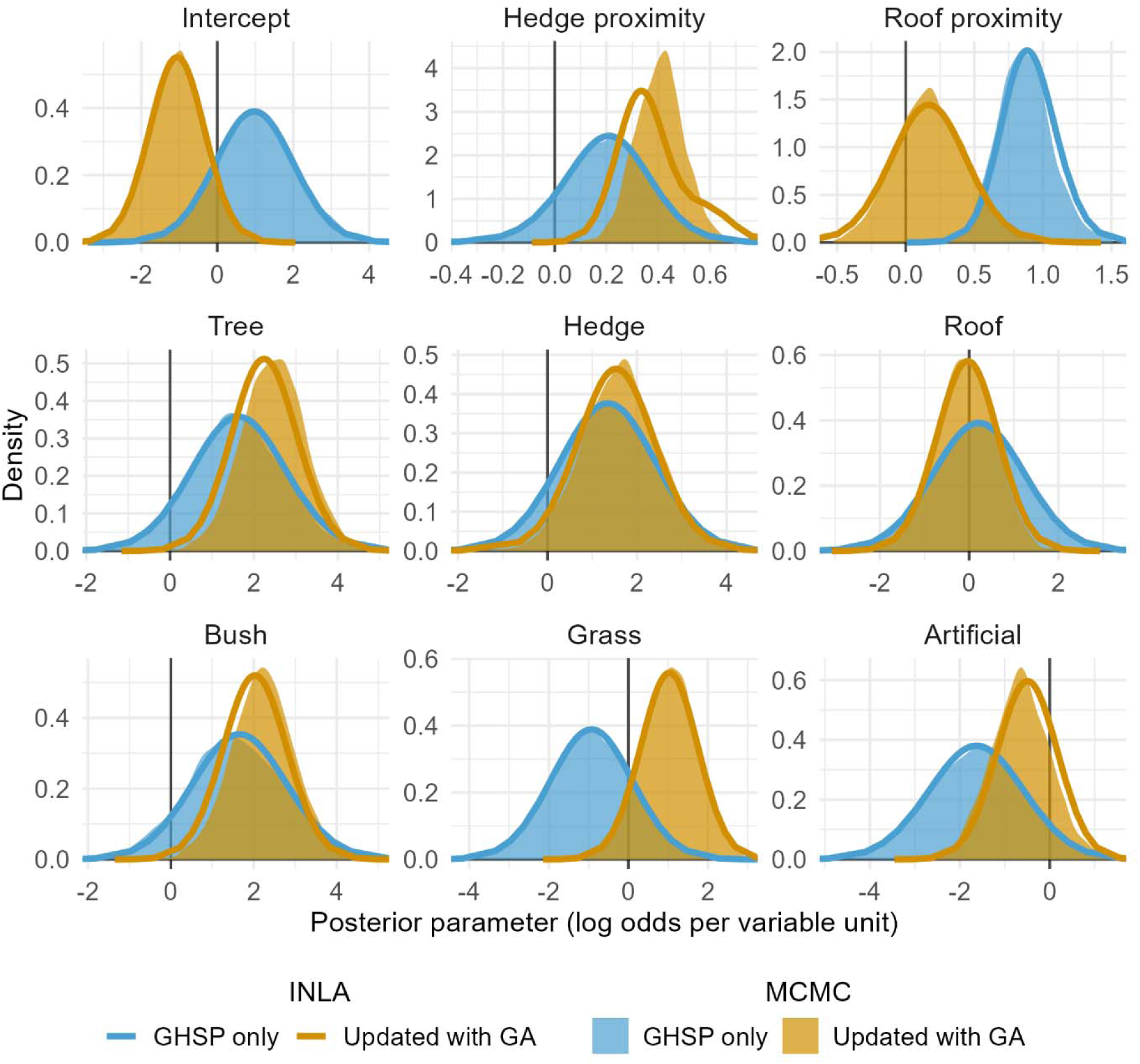
Posterior distributions for model parameters before and after updating of initial GHSP-data model with GA data. Variable units are metres or cell proportion covered, as appropriate. INLA and MCMC fits are both included, to demonstrate that INLA is an acceptable substitute for MCMC in this context.

For roofs and artificial surfaces, on the other hand, the updated model decreased its estimates of impact (positive and negative, respectively). Here, the most salient change was for roof proximity, with a 98% probability that this represents a real difference. This change also implied a reduction in confidence about the effect.

### Effect of model updating on predictive power

Updating with GA data improved the model’s overall ability to predict the GA data, decreased its ability to predict the GHSP data, and increased its ability to predict all data together (Figure 5). Brier decompositions indicated that the improved ability to predict the GA data was the result of improvements in both discrimination and miscalibration. The relevant preregistered hypothesis, of improvements in overall Brier score and its decompositions when predicting GA data, was thus confirmed.^3^ With regard to miscalibration, but not discrimination, the updated model’s overall ability to predict both datasets combined was also improved.

**Figure 5.**
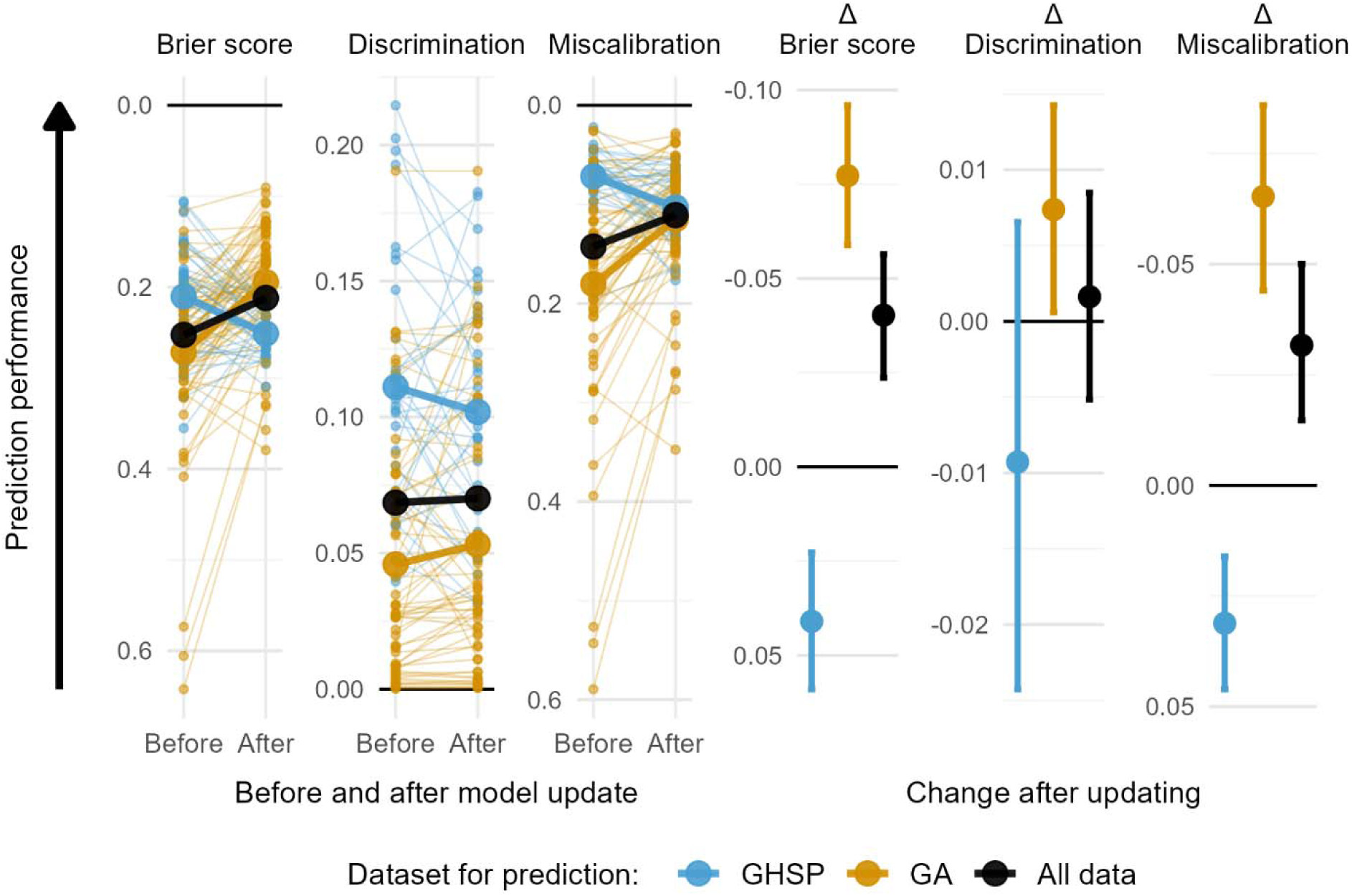
Model performance (leave-one-out cross-validated) before and after updating. The before model is constructed only with GHSP data and the after model is the same model after updating with GA data. Colour indicates dataset for prediction, not dataset for training. Vertical scale direction is set so upwards always represents improvement. For Brier score and decompositions (left hand panels), small points and lines show performance for individual locations (GA gardens and GHSP colonies), while large points and lines show mean performance. Change scores (right-hand panels) are median and 95% credible intervals.

A concern with leave-one-location-out cross-validation relates to the possibility that sparrow-habitat associations, or habitats themselves, vary spatially, with nearer locations more similar than distant ones. This could cause cross-validated predictive performance estimates to be optimistically biased, because locations near to a held-out location remain in the training set (Dormann et al., 2007). Spatial autocorrelation analysis of model residuals indicated no evidence for greater similarity between nearer GA gardens, strongly mitigating this concern, but we nevertheless carried out an additional (non-preregistered) leave-one-out analysis as a precautionary robustness check, dividing the UK into four geographic regions, and leaving one region out at a time. This yielded a near-identical mean Brier score to that from leave-one-location-out (location: 0.19, 95% CI [0.18, 0.21]; region: 0.20, 95% CI [0.18, 0.22]), indicating that the leave-one-location-out estimate was not inflated by spatial dependence between nearby GA gardens, and the concern is therefore allayed. Further details of both these analyses are in Supporting Information Section 4.3.3.1.

For the current EDS application, thresholding model output to produce a presence-absence prediction is unnecessary, but in many contexts, decision making requires a best guess. We therefore explored thresholded presence-absence predictions for GA data, finding at least small improvements for the updated model, depending on threshold selection approach (Supporting Information Section 4.3.3.2).

### Effect consistency

An aspect of model prediction particularly relevant in decision-making contexts is effect consistency, defined as the estimated probability that a random location will display an effect in the predicted direction. This is correlated with but separable from the estimated fixed effect, because it also depends on inter-location effect heterogeneity. Figure 6 illustrates good confidence in most vegetation-related effects and effect consistencies – for example, trees are predicted to have a positive effect in approximately 80% of gardens. There is some weak evidence that the consistency of hedge effects might be slightly greater than expected, given their fixed effect sizes, although trees and bushes are estimated to be most beneficial. The equivalent figure for the GHSP-only model (Supporting Information Section 4.3.4.1) indicates that GHSP data by itself does not enable confident predictions except for those based on roof proximity and artificial surface.

**Figure 6.**
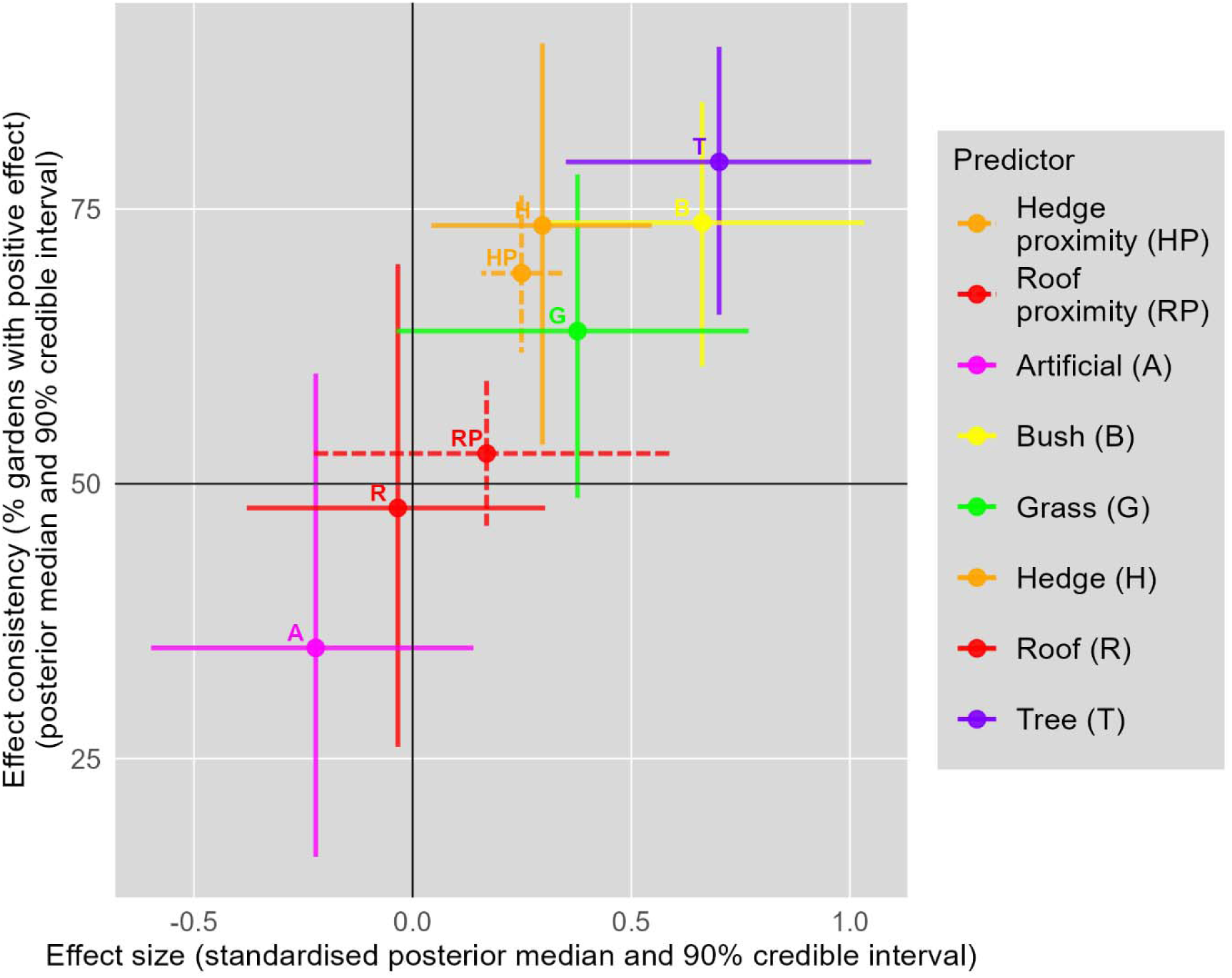
Effect consistency and standardised effect size across predictors for GA-updated MCMC model. Effect sizes are standardized by multiplying regression parameters by the pooled within-location standard deviation for each predictor.

### Decision-support-seeking behaviours

Participants were paid small sums to follow specific instructions within the GA system, necessary to provide data that could support model updating. They were not instructed to obtain environmental decision support by running models to see their garden’s scores for sparrows or carbon. However, 56% examined their sparrow scores, 59% examined their carbon scores, 76% examined at least one of those, 18% planned changes to their garden using planning mode, and 11% examined how these changes affected scores.

## Discussion

### Observed model improvements

The REDS approach was successfully demonstrated in a proof-of-concept system, Garden Advice (GA). Users contributed habitat and species occurrence data, refining the decision support available to later users by increasing the accuracy of the model that generated the predictions. The improved performance when predicting the GA data obtained from across the UK implies more accurate predictions regarding the general UK sparrow population. Under the REDS approach, the population from which data is obtained to update the model is automatically the same population the system is designed to support, which is not always the case with EDS.

There was good confidence in improvements in the model’s abilities to utilise habitat type to predict sparrow presence (discrimination) and to predict overall prevalence (calibration). The measured improvement in discrimination was modest, so the overall improvement was due mainly to improved calibration, although the change score credible intervals (Figure 5, right-hand panels) indicate the discrimination improvement was also real.

Improved discrimination is an expected outcome of posteriors that are narrower and shifted, reflecting improved confidence in modified estimates of the habitat associations. Predictive performance was reduced for the original GHSP data set, which can be explained as follows. Although we did not find evidence for spatial variation in habitat association patterns across the UK, it is highly unlikely that there are no undetected effects. Any such effects would imply that predictive performance for data from one area (the GHSP data is from Glasgow) would reduce when the model is updated with data from across the UK.

The Brier score for predicting GA data improved from 0.27 to 0.19 after model updating (lower is better), both of which are poor scores in the context, suggesting noisy data that limits the achievable performance of predictive modelling. One likely reason is that we included no global covariates describing garden context, such as surrounding habitat, physical geography, or predators. Such variables are possible to obtain through the current CS approach, or from other databases, but could not be included in the current study due to their absence in the GHSP data set.

### Changes to specific model parameters after model updating

Whether the changes to posteriors reflect reality depends on the veracity of the contributed data, but literature suggests that the most marked parameter change represents real ecological information. Weak confidence in a negative association between grass and sparrow occurrence updated to weak confidence in a positive association. Consistent evidence includes anecdotal (Bradbury, 2024; Teo, 2023) and systematic (Vincent, 2005) observations of House Sparrows indicating that although grass is not a preferred foraging habitat, they can obtain seeds and invertebrates there. In Spain, reduction of grass in urban parks has been associated with House Sparrow decline (Bernat-Ponce et al., 2020).

Other studies indicate negative associations with grass, however (Havlíček, 2021; Weir, 2015). One likely resolution to this conflicting evidence relates to temporal variation. House Sparrow preference for grass has been observed to fluctuate month-on-month and year-on-year (Vincent, 2005), so grass may be preferred when temporary conditions increase its relative benefits. REDS systems are well suited to provide data over time and across circumstances to support a richer model that could address issues of seasonality and other reasons for heterogenous effects.

Of the remaining changes in model parameters, most represented increased confidence in slightly larger positive effects. This was the case for trees, bushes, and hedges (cover and proximity), and this is also supported by the general literature. There were weak indications that the effects of hedges may be more consistent across locations than would be predicted by effect size alone, according with previous observations of the great importance of hedgerows for birds and reinforcing the importance of conserving hedges (Glasgow House Sparrow Project, n.d.; Hinsley & Bellamy, 2000). Such effect consistency estimates are only available when sampling large numbers of independent locations, as they represent inter-location effect heterogeneity. REDS and other citizen science systems are well suited to collecting such data (Garretson et al., 2023). Effect consistency is important for EDS as it is appropriate for decision support to emphasise consistent effects, independently of effect size, because consistent effects are robust to local idiosyncrasies. We are unaware of previous uses of our specific method for intuitively presenting effect heterogeneity, by transforming to the percentage of locations where the effect is expected to apply. This presentation method could be generally beneficial for decision support.

The initial GHSP-only model’s strong confidence in the association of sparrows with roof proximity was removed by updating the model with GA data. This is unlikely to be realistic, given the close affiliation of House Sparrows with human dwellings (Shaw et al., 2008). This discrepancy highlights a key issue: the accuracy of GA user observations was not evaluated, and the result likely reflects observation bias. Birds on or near roofs are harder to see, especially for those observing from indoors.

### Challenges associated with the automatic incorporation of Citizen Science data

Another study that solicited non-expert animal observations from UK gardens found that although participants could provide accurate habitat descriptions, hedgehog presence-absence reports were uncorrelated with objective data (Williams et al., 2015). Hedgehogs are harder than sparrows for humans to detect, but this nevertheless provides a sobering reminder of the necessity for independent confirmation of citizen science data. However, reviews indicate that citizen science data is often reliable, and measures such as additional training provide benefits (Aceves-Bueno et al., 2017; Backstrom et al., 2024; Johnston et al., 2023; Kosmala et al., 2016). Further, although our study involved data solicitation from a wide public, in practice EDS systems are often used by land managers such as farmers, who might provide more accurate observations. Nevertheless, the next stage in REDS concept validation must involve independent verification of data quality. Such model validation is one of the factors most associated with EDS system uptake (Walling & Vaneeckhaute, 2020).

With REDS systems where data for the same locality is obtained from multiple users or where ground-truth or other independent measures allow estimates of data-contributor reliability, it will be possible to include terms for observation bias in models (Johnston et al., 2018; Kaurila et al., 2026; Santos-Fernandez & Mengersen, 2021). The issue is particularly acute with REDS systems because they will benefit from deployment to a wide audience of users whose data updates the system automatically. It is even possible that users could behave maliciously. In the current study, although model update was automatic, there was manual checking that data contributions plausibly represented UK domestic gardens, resulting in some exclusions. Such checking will be necessary in many cases, although some checks could themselves be automated. We do not recommend wide-scale deployment of REDS systems without careful consideration of this issue.

### Broader implications of REDS in the context of existing citizen science (CS), environmental decision support (EDS), and digital twin (DT) tools

EDS systems predict the effects of potential changes to local conditions so that management can be planned. However, standard EDS relies on static models that may not optimally reflect the reality experienced by current users. There are many reasons why relations between environmental variables may not be static, because they may be moderated by additional variables which are themselves changing (such as climate, Bueno de Mesquita et al., 2021). REDS systems can better remain up to date, in essence informing users “if you change your conditions in such a way, your situation will more closely resemble the situation recently reported by certain other users, so your observations will become more like theirs.” A concrete example from this study is that, thanks to the provision of user data that updated the model to separate grass from artificial surfaces, by the end of the study period the system advised that lawns are better for sparrows than decking or paving. Although our users arrived at the system because of payment, and were never instructed to use the EDS functions, most of them did request model predictions. This means that later users did benefit from the improvements to advice generated by earlier data contributions.

Future REDS systems would benefit from more complex models. Habitat association models do not consider causal mechanisms and do not necessarily predict long-term species health, because local observations may not predict longer term demographic changes (Bacon et al., 2017; Matthiopoulos et al., 2019; Regos et al., 2021). That said, reviews indicate that although there are important exceptions, the availability of preferred habitats does tend to predict species abundance (Boyce et al., 2016; Weber et al., 2017). In principle, the REDS citizen science approach could also track demographic change, as one route to support more complex and accurate models.

As reviewed in the Introduction, neither CS nor DT systems are inherently REDS, and we found no such existing systems that utilise user data to update models for predicting consequences of potential management decision, which is the defining feature of REDS. Many such systems are close, however. Consider eBird, for example, which does update thanks to user input, and already aids management decisions by informing where birds are (Farwell et al., 2026). With the right models, it could further inform users, by explicitly modelling the predicted impact of management decisions.

Some DT systems are informed by user input, and some are intended to provide predictive models for management decisions – but we are unaware of any that yet do both, despite recent reviews (Hazeleger et al., 2024; Trantas et al., 2023). The added complexity of DTs, involving synchronous modelling of a specific physical system, may be one reason – the reviews indicate appreciable challenges in the development of DTs. Simpler habitat-association models may sometimes suffice, although it seems likely that some DTs may soon become REDS.

Further development of the REDS concept will benefit from closer examination of the potential social synergies generated by combining CS and EDS. There is always potential for useful synergy when closing an information loop, allowing feedback mechanisms to operate (Wiener, 1948). For REDS, some of these synergies might relate to the strong human motivation towards reciprocity (Bowles & Gintis, 2011). It is likely that public data sharing that already exists on platforms such as iNaturalist and PurpleAir is motivated in large part in this way (Metzger & Runge, 2023; Pai & Tsai, 2016; Truong & Van der Wal, 2024).

Examples of existing initiatives that could become REDS include membership-organisation CS initiatives, such as the British Trust for Ornithology’s Garden Birdwatch (Morrison et al., 2014), schemes associated with the Bumblebee Conservation Trust (Sharma et al., 2019), or the Game & Wildlife Conservation Trust’s Partridge Count Scheme (Ewald et al., 2009). Participation could be increased, and relevant environmental management behaviour could be improved, as citizens receive tailored advice as an immediate and integral part of reporting their contributions.

Additional motivation for engagement may come from participants more directly seeing how individual contributions improve centralised knowledge. Stakeholder engagement is a key determinant of EDS system success (Walling & Vaneeckhaute, 2020). Provided that careful consideration is given to the structure of the initial model, REDS can save time for scientists operating such schemes, as analysing and reporting the data is an automated and integrated part of the data-collection process itself. Like DTs, the REDS concept is at an early stage of development, and the next stages must focus on how development and deployment can be achieved without sacrificing the quality of analysis and conclusions.

In summary, the sense that one is helping at the same time as being helped could be a strong factor for further activating the synergies between EDS and CS that this study indicates REDS can provide. These synergies offer positive feedback loops potentially generating a virtuous cycle of greater user engagement, data provision, advice quality, and advice uptake.

## Data availability statement

Data is available via Supporting Information.

## Acknowledgements

Thanks to Nicholas Aebischer and Johan Vegelius for useful discussions, to Prolific-recruited participants who went the extra mile, and to Bridget Kenward for much practical assistance. This study was funded by the EU Horizon grant PRO-COAST (Project ID 101082327).

## Author contributions

Conceptualisation: BK, NC, JE, RK; Data curation: BK, NC; Analysis: BK; Funding acquisition: JE, RK; Methodology: all authors; Software: NC, BK; Writing (original draft): BK; Writing (review): all authors.

## Footnotes

1 Non-predictive databases can also support environmental decisions, but in line with previous literature requiring EDS systems to feature “analysis” or “models” or to be “intelligent” (McIntosh et al., 2011), we exclude such systems from our definition.

2 The probability that a point from the first posterior differs from a point in the second posterior in the same direction as the observed difference in medians.

3 There was one additional preregistered hypothesis, that the updated model would explain variance; this hypothesis is ignored as it is trivial given that the primary hypothesis is confirmed.

## References

1. Aceves-Bueno, E., Adeleye, A. S., Feraud, M., Huang, Y., Tao, M., Yang, Y., & Anderson, S. E. (2017). The Accuracy of Citizen Science Data: A Quantitative Review. The Bulletin of the Ecological Society of America, 98(4), 278–290.

2. Albert, D. A., & Smilek, D. (2023). Comparing attentional disengagement between Prolific and MTurk samples. Scientific Reports, 13(1), 20574.

3. Backstrom, L. J., Callaghan, C. T., Leseberg, N. P., Sanderson, C., Fuller, R. A., & Watson, J. E. M. (2024). Assessing adequacy of citizen science datasets for biodiversity monitoring. Ecology and Evolution, 14(2), e10857.

4. Bacon, L., Hingrat, Y., Jiguet, F., Monnet, A.-C., Sarrazin, F., & Robert, A. (2017). Habitat suitability and demography, a time-dependent relationship. Ecology and Evolution, 7(7), 2214–2222.

5. Bernat-Ponce, E., Gil-Delgado, J. A., & López-Iborra, G. M. (2020). Replacement of semi-natural cover with artificial substrates in urban parks causes a decline of house sparrows Passer domesticus in Mediterranean towns. Urban Ecosystems, 23(3), 471–481.

6. Bondia-Barceló, J., Castellà-Roca, J., & Viejo, A. (2016). Building Privacy-Preserving Search Engine Query Logs for Data Monetization. 2016 Intl IEEE Conferences on Ubiquitous Intelligence & Computing, Advanced and Trusted Computing, Scalable Computing and Communications, Cloud and Big Data Computing, Internet of People, and Smart World Congress (UIC/ATC/ScalCom/CBDCom/IoP/SmartWorld),

7. Bowles, S., & Gintis, H. (2011). A cooperative species: Human reciprocity and its evolution. Princeton University Press.

8. Boyce, M. S., Johnson, C. J., Merrill, E. H., Nielsen, S. E., Solberg, E. J., & van Moorter, B. (2016). REVIEW: Can habitat selection predict abundance? Journal of Animal Ecology, 85(1), 11–20.

9. Bradbury, K. (2024). The house sparrow (Passer domesticus). https://www.encounter-nature.com/features/the-house-sparrow-passer-domesticus

10. Bressler, R. D. (2021). The mortality cost of carbon. Nature Communications, 12(1), 4467.

11. Bueno de Mesquita, C. P., White, C. T., Farrer, E. C., Hallett, L. M., & Suding, K. N. (2021). Taking climate change into account: Non-stationarity in climate drivers of ecological response. Journal of Ecology, 109(3), 1491–1500.

12. Burris, M., Khan, M., & Johnson, J. (2023). “Is That Route Really the Most Fuel-Efficient?” (National Institute For Congestion Reduction Research Reports, Issue 17). 10.5038/CUTR-NICR-Y3-4-8

13. Cole, E., Van Horn, G., Lange, C., Shepard, A., Leary, P., Perona, P., Loarie, S., & Mac Aodha, O. (2023). *Spatial Implicit Neural Representations for Global-Scale Species Mapping* Proceedings of the 40th International Conference on Machine Learning, Proceedings of Machine Learning Research. https://proceedings.mlr.press/v202/cole23a.html

14. Cool Farm. (2025a). Cool Farm. Retrieved 2025-10-03 from https://coolfarm.org/

15. Cool Farm. (2025b). Cool Farm: Frequently Asked Questions. Retrieved 2025-10-03 from https://coolfarm.org/frequently-asked-questions/

16. De Antoni, A., Rucco, M., Cattaneo, A. M., Gezer, E., Sulis, G., Draicchio, P., Iacca, G., Pugliese, A., & Mancini, M. V. (2026). PesTwin: a biology-informed Digital Twin for enabling precision farming. *arXiv preprint, arXiv:2603.12294*.

17. De Koning, K. (2025). The crane radar: Development and deployment of an operational eco-digital twin. Ecological Informatics, 85, 102938.

18. de Koning, K., Broekhuijsen, J., Kühn, I., Ovaskainen, O., Taubert, F., Endresen, D., Schigel, D., & Grimm, V. (2023). Digital twins: dynamic model-data fusion for ecology. Trends in Ecology & Evolution, 38(10), 916–926.

19. De Laet, J., & Summers-Smith, J. D. (2007). The status of the urban house sparrow Passer domesticus in north-western Europe: a review. Journal of Ornithology, 148(2), 275–278.

20. Dormann, C. F., McPherson, J. M., Araújo, M. B., Bivand, R., Bolliger, J., Carl, G., Davies, R. G., Hirzel, A., Jetz, W., Daniel Kissling, W., Kühn, I., Ohlemüller, R., Peres-Neto, P. R., Reineking, B., Schröder, B., Schurr, F. M., & Wilson, R. (2007). Methods to account for spatial autocorrelation in the analysis of species distributional data: a review. Ecography, 30(5), 609–628.

21. Ewald, J. A., Kingdon, N. G., & Santin-Janin, H. (2009). The GWCT Partridge Count Scheme: a volunteer-based monitoring and conservation promotion scheme. In S. B. Cederbaum, B. C. Faircloth, T. M. Terhune, J. J. Thompson, & J. P. Carroll (Eds.), Gamebird 2006: Quail VI and Perdix XII (pp. 27–37). Warnell School of Forestry and Natural Resources.

22. Farwell, L. S., Stillman, A. N., Lancaster, J. D., Turbek, S. P., Muñoz, J. M., Peele, A. M., Rylander, R. J., Yanega, G. M., Iglecia, M. N., Robinson, O. J., Duren, A. M., Hannuksela, A., Huang, A., Yarris, G. S., & Eger, C. G. (2026). Leveraging eBird data products to inform regional bird conservation priorities and objectives. Ornithological Applications, 128(1), 1–14.

23. Fraisl, D., Hager, G., Bedessem, B., Gold, M., Hsing, P.-Y., Danielsen, F., Hitchcock, C. B., Hulbert, J. M., Piera, J., Spiers, H., Thiel, M., & Haklay, M. (2022). Citizen science in environmental and ecological sciences. Nature Reviews Methods Primers, 2(1), 64.

24. Garretson, A., Cuddy, T., Duffy, A. G., iNaturalist Citizen, S., & Forkner, R. E. (2023). Citizen science data reveal regional heterogeneity in phenological response to climate in the large milkweed bug, Oncopeltus fasciatus. Ecology and Evolution, 13(7), e10213.

25. Ginsberg, J., Mohebbi, M. H., Patel, R. S., Brammer, L., Smolinski, M. S., & Brilliant, L. (2009). Detecting influenza epidemics using search engine query data. Nature, 457(7232), 1012–1014.

26. Glasgow House Sparrow Project. (n.d.). Help your sparrows. Retrieved 2025-12-15 from https://www.housesparrowscience.com/help-your-sparrows/

27. Hanson, H. E., Mathews, N. S., Hauber, M. E., & Martin, L. B. (2020). The house sparrow in the service of basic and applied biology. Elife, 9.

28. Havlíček, J. (2021). The breeding and foraging ecology of the House Sparrow in rural and urban environments [PhD Thesis, University of South Bohemia in České Budějovice].

29. Hazeleger, W., Aerts, J. P. M., Bauer, P., Bierkens, M. F. P., Camps-Valls, G., Dekker, M. M., Doblas-Reyes, F. J., Eyring, V., Finkenauer, C., Grundner, A., Hachinger, S., Hall, D. M., Hartmann, T., Iglesias-Suarez, F., Janssens, M., Jones, E. R., Kölling, T., Lees, M., Lhermitte, S., . . . Vossepoel, F. C. (2024). Digital twins of the Earth with and for humans. Communications Earth & Environment, 5(1), 463.

30. Hinsley, S. A., & Bellamy, P. E. (2000). The influence of hedge structure, management and landscape context on the value of hedgerows to birds: A review. Journal of Environmental Management, 60(1), 33–49.

31. Johnston, A., Fink, D., Hochachka, W. M., & Kelling, S. (2018). Estimates of observer expertise improve species distributions from citizen science data. Methods in Ecology and Evolution, 9(1), 88–97.

32. Johnston, A., Matechou, E., & Dennis, E. B. (2023). Outstanding challenges and future directions for biodiversity monitoring using citizen science data. Methods in Ecology and Evolution, 14(1), 103–116.

33. Johnston, A., Rodewald, A. D., Strimas-Mackey, M., Auer, T., Hochachka, W. M., Stillman, A. N., Davis, C. L., Ruiz-Gutierrez, V., Dokter, A. M., Miller, E. T., Robinson, O., Ligocki, S., Jaromczyk, L. O., Crowley, C., Wood, C. L., & Fink, D. (2025). North American bird declines are greatest where species are most abundant. Science, 388(6746), 532–537.

34. Kaurila, K., Kuningas, S., Lappalainen, A., & Vanhatalo, J. (2026). Species distribution modeling with expert elicitation and Bayesian calibration. Ecography, 2026(3), e08173.

35. Kayatz, B., Baroni, G., Hillier, J., Lüdtke, S., Heathcote, R., Malin, D., van Tonder, C., Kuster, B., Freese, D., Hüttl, R., & Wattenbach, M. (2019). Cool Farm Tool Water: A global on-line tool to assess water use in crop production. Journal of Cleaner Production, 207, 1163–1179.

36. Kenward, B. (2024). The 1000-ton rule, which emphasises the mortality cost of carbon, can be an effective persuasion tool, including for use by activists. https://www.benkenward.com/articles/the_1000_ton_rule_can_be_an_effective_persuasion_tool.pdf

37. Kenward, R., Papathanasiou, J., Manos, B., & Arampatzis, E. (Eds.). (2013). Transactional Environmental Support System Design: Global Solutions. IGI Global.

38. Kosmala, M., Wiggins, A., Swanson, A., & Simmons, B. (2016). Assessing data quality in citizen science. Frontiers in Ecology and the Environment, 14(10), 551–560.

39. Lu, B., Francescutto, L., Howie, S., Lin, H., Wu, Q., Hedley, N., Jamali, A., & McDonald, I. (2026). Exploring the concept of digital twins of wetlands for supporting ecosystem monitoring and management. Big Earth Data, 10(1), 37–67.

40. Matthiopoulos, J., Field, C., & MacLeod, R. (2019). Predicting population change from models based on habitat availability and utilization. Proceedings of the Royal Society B: Biological Sciences, 286(1901), 20182911.

41. McIntosh, B. S., Ascough, J. C., Twery, M., Chew, J., Elmahdi, A., Haase, D., Harou, J. J., Hepting, D., Cuddy, S., Jakeman, A. J., Chen, S., Kassahun, A., Lautenbach, S., Matthews, K., Merritt, W., Quinn, N. W. T., Rodriguez-Roda, I., Sieber, S., Stavenga, M., . . . Voinov, A. (2011). Environmental decision support systems (EDSS) development – Challenges and best practices. Environmental Modelling & Software, 26(12), 1389–1402.

42. McKinley, D. C., Miller-Rushing, A. J., Ballard, H. L., Bonney, R., Brown, H., Cook-Patton, S. C., Evans, D. M., French, R. A., Parrish, J. K., Phillips, T. B., Ryan, S. F., Shanley, L. A., Shirk, J. L., Stepenuck, K. F., Weltzin, J. F., Wiggins, A., Boyle, O. D., Briggs, R. D., Chapin, S. F., . . . Soukup, M. A. (2017). Citizen science can improve conservation science, natural resource management, and environmental protection. Biological Conservation, 208, 15–28.

43. Merkle, E. C., & Hartman, R. (2018). Weighted Brier score decompositions for topically heterogenous forecasting tournaments. Judgment and Decision Making, 13(2), 185–201.

44. Metzger, F. M., & Runge, G. (2023). Reciprocity of Data Sharing Infrastructures: A Conceptual Norms Framework. In A. Roßnagel, M. Friedewald, C. L. Geminn, M. Karaboga, & S. Schindler (Eds.), Data Sharing – Datenkapitalismus by Default? 10.24406/publica-2072

45. Morrison, C. A., Robinson, R. A., Leech, D. I., Dadam, D., & Toms, M. P. (2014). Using citizen science to investigate the role of productivity in House Sparrow Passer domesticus population trends. Bird Study, 61(1), 91–100.

46. Nghiem, L. T. P., Papworth, S. K., Lim, F. K. S., & Carrasco, L. R. (2016). Analysis of the Capacity of Google Trends to Measure Interest in Conservation Topics and the Role of Online News. PLOS ONE, 11(3), e0152802.

47. ONS Geography. (2025). Rural Urban Classification (2021) of Output Areas in EW. Retrieved 2025-05-19 from https://geoportal.statistics.gov.uk/datasets/ons::rural-urban-classification-2021-of-output-areas-in-ew/about

48. Ovaskainen, O., Winter, S., Tikhonov, G., Lauha, P., Lehtiö, A., Nokelainen, O., Abrego, N., Aroluoma, A., Harrison, J. P., Heikkinen, M., Kallio, A., Koliseva, A., Lehikoinen, A., Roslin, T., Somervuo, P., Souza, A. T., Tahir, J., Talaskivi, J., Turunen, A., . . . Dunson, D. (2026). A digital twin for real-time biodiversity forecasting with citizen science data. Nature Ecology & Evolution, 10(3), 481–495.

49. Pai, P., & Tsai, H.-T. (2016). Reciprocity norms and information-sharing behavior in online consumption communities: An empirical investigation of antecedents and moderators. Information & Management, 53(1), 38–52.

50. Papathanasiou, J., & Kenward, R. (2014). Design of a data-driven environmental decision support system and testing of stakeholder data-collection. Environmental Modelling & Software, 55, 92–106.

51. Parker, T., Fraser, H., & Nakagawa, S. (2019). Making conservation science more reliable with preregistration and registered reports. Conservation Biology, 33(4), 747–750.

52. Pearce, J. M., & Parncutt, R. (2023). Quantifying Global Greenhouse Gas Emissions in Human Deaths to Guide Energy Policy. Energies, 16(16).

53. Peer, E., Rothschild, D., Gordon, A., Evernden, Z., & Damer, E. (2022). Data quality of platforms and panels for online behavioral research. Behavior Research Methods, 54(4), 1643–1662.

54. PurpleAir. (2025). PurpleAir. Retrieved 2025-10-03 from https://purpleair.com/

55. Regos, A., Tapia, L., Gil-Carrera, A., & Domínguez, J. (2021). Caution Is Needed When Using Niche Models to Infer Changes in Species Abundance: The Case of Two Sympatric Raptor Populations. Animals, 11(7), 2020.

56. RHS. (2025). RHS State of Gardening Report 2025. https://www.rhs.org.uk/about-us/what-we-do/rhs-state-of-gardening

57. Robinson, D. L., Goodman, N., & Vardoulakis, S. (2023). Five Years of Accurate PM2.5 Measurements Demonstrate the Value of Low-Cost PurpleAir Monitors in Areas Affected by Woodsmoke. International Journal of Environmental Research and Public Health, 20(23).

58. Sandhaus, S., Kaufmann, D., & Ramirez-Andreotta, M. (2019). Public participation, trust and data sharing: gardens as hubs for citizen science and environmental health literacy efforts. International Journal of Science Education, Part B, 9(1), 54–71.

59. Santos-Fernandez, E., & Mengersen, K. (2021). Understanding the reliability of citizen science observational data using item response models. Methods in Ecology and Evolution, 12(8), 1533–1548.

60. Sharma, N., Greaves, S., Siddharthan, A., Anderson, H. B., Robinson, A.-M., Colucci-Gray, L., Wibowo, A. T., Bostock, H., Salisbury, A., & Roberts, S. (2019). From citizen science to citizen action: analysing the potential for a digital platform to cultivate attachments to nature. Journal of Science Communication, 18(01), A07.

61. Shaw, L. M., Chamberlain, D., & Evans, M. (2008). The House Sparrow Passer domesticus in urban areas: reviewing a possible link between post-decline distribution and human socioeconomic status. Journal of Ornithology, 149(3), 293–299.

62. Siegert, S. (2017). Simplifying and generalising Murphy’s Brier score decomposition. Quarterly Journal of the Royal Meteorological Society, 143(703), 1178–1183.

63. Taubert, F., Rossi, T., Wohner, C., Venier, S., Martinovič, T., Khan, T. H., Gordillo, J. L., & Banitz, T. (2024). Prototype Biodiversity Digital Twin: grassland biodiversity dynamics [10.3897/rio.10.e124168]. Research Ideas and Outcomes, 10, e124168.

64. Teo, L. W. (2023). House Sparrows dig and chew on grass stolons. https://besgroup.org/2023/05/18/house-sparrows-dig-chew-grass-stolons/

65. Trantas, A., Plug, R., Pileggi, P., & Lazovik, E. (2023). Digital twin challenges in biodiversity modelling. Ecological Informatics, 78, 102357.

66. Truong, M.-X. A., & Van der Wal, R. (2024). Exploring the landscape of automated species identification apps: Development, promise, and user appraisal. BioScience, 74(9), 601–613.

67. Vehtari, A. (2025). Can cross-validation be used for hierarchical / multilevel models? https://users.aalto.fi/~ave/CV-FAQ.html#8_Can_cross-validation_be_used_for_hierarchicalmultilevel_models

68. Vincent, K. (2005). Investigating the causes of the decline of the urban house sparrow Passer domesticus in Britain [PhD Thesis, De Montfort University, Leicester].

69. Walling, E., & Vaneeckhaute, C. (2020). Developing successful environmental decision support systems: Challenges and best practices. Journal of Environmental Management, 264, 110513.

70. Weber, M. M., Stevens, R. D., Diniz-Filho, J. A. F., & Grelle, C. E. V. (2017). Is there a correlation between abundance and environmental suitability derived from ecological niche modelling? A meta-analysis. Ecography, 40(7), 817–828.

71. Weir, J. E. S. (2015). Urban green space management for invertebrates and house sparrows [PhD Thesis, Imperial College London, Department of Ecology and Evolution].

72. Wiener, N. (1948). Cybernetics: Or Control and Communication in the Animal and the Machine. MIT University Press.

73. Williams, R. L., Stafford, R., & Goodenough, A. E. (2015). Biodiversity in urban gardens: Assessing the accuracy of citizen science data on garden hedgehogs. Urban Ecosystems, 18(3), 819–833.

74. Wong-Parodi, G., Mach, K. J., Jagannathan, K., & Sjostrom, K. D. (2020). Insights for developing effective decision support tools for environmental sustainability. Current Opinion in Environmental Sustainability, 42, 52–59.

