## Supporting Information for "Reciprocal Environmental Decision Support (REDS): better tailored advice in return for data"

### *2. Document overview*

This Supporting Information document contains all information or links necessary to fully describe the study. The individual sections are as follows:

*Document overview*: The current section.

*Preregistration*: Link for the study preregistration and related information.

*Supporting Information for Main Text*: Further details on matters described more briefly in the main text, structured with the same headers as the main text for ease of navigation.

*Data and Code Package*: Link and description for an online repository containing code and data used for model updating within Garden Advice and subsequent analysis of the data presented in the main text.

*Garden Advice App (REDS_ONE) Technical Architecture Description*: An overview of the architecture of the web app used to collect data.

*Garden Advice App (REDS_ONE) Code Package*: Link for an online repository containing code that can be used to understand further details of the app’s behaviour.

*Garden Advice Carbon Model*: Describes the Garden Advice carbon model.

*Recruitment Materials*: Two documents used as part of recruitment – the study advertisement within Prolific, and the screening questionnaire.

*References*: References for sources cited in Supporting Information.

### *3. Preregistration*

#### 3.1. Link and description

The preregistration is available at the following link [BLINDED FOR PEER-REVIEW]:

https://osf.io/xkeqp/overview?view_only=d8191b3752a24e639a47e7510cec763e

The primary preregistration document contains preregistered hypotheses and methods that are also included in the main text or in this Supporting Information document and are therefore not required for understanding the study. All analytical methods reported are as per the preregistration or are explicitly marked as exploratory or as deviations from preregistration (see list below). Ecologists unfamiliar with the rationale for preregistration are encouraged to consult Parker et al. (2019).

Additional preregistered material represents exploration of original code provided by Matthiopoulos et al. (2019), and subsequent pilot modelling work, showing how their original model was explored and then changed to provide the initial model for current purposes. That material may be of limited interest, because the adaptation resulted in an initial model that is appreciably different to Matthiopoulos et al.’s own model (2019).

#### 3.2. List of all deviations from preregistration

1. H1 was not tested, because it is trivial given that H2 (the primary hypothesis) is confirmed.

2. Increasing the conservativeness of our comparison of the initial and updated models, we adjust the initial model offset to use the final prevalence information (see main text Methods sub-section “Model predictions and their evaluation”).
3. We use INLA instead of MCMC to fit models for cross-validated comparison (see main text Methods sub-section “Model specification and updating” and corresponding sub-section below).
4. We used non-default priors for the INLA random slope hyperparameters (see main text Methods sub-section “Model specification and updating” and corresponding sub-section below).
5. We used different code to that preregistered to calculate Brier scores and their decompositions (see “Brier score calculations” sub-section below).

### *4. Supporting Information for Main Text*

#### 4.1. Introduction

##### *4.1.1. Environmental decision support*

###### 4.1.1.1. The systematic search for other REDS systems

The systematic search for existing REDS systems was intended to be exhaustive enough to make it likely to find other REDS tools unless they are very rare or non-existent. A more exhaustive search would be extremely challenging, given the large number of existing EDS tools. Through general internet searches and pre-existing knowledge, we selected eight different lists and searches and identified 241 tools for assessment (Table S1). The database of these tools and a script to summarise them are contained in the Data and Code Package.

The 241 recorded tools were all those considered relevant after cursory examination of all items in the lists and search results. Reasons an item was not considered relevant included that it was not software, that information was not available, or that it was clearly not related to environmental management. Of these 241 listed tools, 7 could not be found. Assessment of the remaining 234 was conducted by finding the tool itself or a webpage or scientific article describing it, and coding properties as present, absent, or unknown (Table S2). It is possible that unknown was sometimes coded despite available relevant information because it could not be found in the time available for coding.

Table S1. List of lists used to systematically collect tools.

| List or search | Identified tools |
|---|---|
| List of decision support systems for natural hazard risk reduction in Appendix B of Newman et al. (2017). | 60 |
| Three Web of Science searches aimed at finding applications similar to Garden Advice: 1. (participatory OR "citizen science" OR crowdsourced) AND (app OR application OR tool OR platform OR software) AND (biodiversity OR "carbon storage" OR "carbon sequestration") – 2015-2025; 2. ("participatory modelling" OR "citizen science" OR "user-contributed data" OR crowdsourced) AND ("model updating" OR "adaptive model" OR "model calibration" OR "feedback loop" OR "iterative model improvement") – any time; 3. garden AND (app OR application OR tool OR platform OR software) AND (biodiversity OR "carbon storage" OR "carbon sequestration" – any time. | 57 |
| List of models maintained by the Hydrologic Modelling Community of Practice (https://www.epa.gov/hydrowq). | 37 |
| List of "Agri-Environmental Decision Support System Tools" maintained by the Ontario Federation of Agriculture (https://ofa.on.ca/resources/agri-environmental-decision-support-system-tools/). | 35 |
| List of "Data and Tools" maintained by the US Department of Agriculture Forest Service (https://research.fs.usda.gov/products/dataandtools), filtering by Type = "Decision Support System" or "Model". | 16 |
| List of "Policy Support Tools" maintained by IPBES (https://www.ipbes.net/policy-support/search), filtering by Option = "Policy Support Tool". | 13 |
| List of "Tools and Data" maintained by the US Climate Resilience Toolkit (https://toolkit.climate.gov/tool), filtering by Step = "Investigate options" and (separately) by Search = "Model". | 12 |
| Non-systematic pre-existing knowledge | 6 |
| List of "Climate Change Decision Support Tools" maintained by UK Forest Research (https://www.forestresearch.gov.uk/climate-change/resources/decision-support-tools/) | 5 |

Table S2. REDS-relevant properties coded as present or not present in tools.

| Property | Property known (% of all tools) | Property present (% of known) |
|---|---|---|
| Operates over a network | 91% | 60% |
| Makes model predictions available to user | 88% | 72% |
| Is EDS (model predictions support environmental management) | 78% | 41% |
| Model updated by user-contributed data (reciprocal) | 83% | 3% |
| Model predictions based on environmental condition variables | 88% | 56% |

Reciprocal updating (where user input to a system is used to update predictive models) is evidently rare in environmental tools. The 3% of assessed tools with this property correspond to five tools: Artsorakel (Naturalis Biodiversity Centre, 2020), iNaturalist (Mason et al., 2025), eBird (Gorleri et al., 2023), Pl@ntNet (Lefort et al., 2024), Merlin Bird ID (Cornell Lab of Ornithology, 2025), and the Crane Radar (De Koning, 2025). The first four are all identification apps and have been coded as reciprocal because photographs or sound recordings submitted together with identifications can be used to train the species identification model. The Crane Radar is a digital twin allowing real-time predictions about the locations of Common Crane *Grus*

*grus* migratory groups – observations from birdwatchers are a key part of the input. iNaturalist has additional reciprocal functionality as described in the main text. Importantly, none of these tools was classed as an Environmental Decision Support tool, because they are not designed with the primary function of directly supporting environmental management decisions. In line with this, none of them except the Crane Radar uses environmental conditions predictor variables.

#### 4.2. Methods

##### *4.2.1. Declarations*

For details of LLM use in writing of code see top-level sections "Data and Code Package" and "Garden Advice App (REDS_ONE) Code Package". For details of informed consent collection, see "Recruitment" subsection and top-level section "Recruitment materials". For details of preregistration, see relevant top-level section.

##### *4.2.2. Participants*

###### 4.2.2.1. Recruitment

In early pilots, users were recruited via social media advertisements emphasising advice from the carbon model; recruitment for later pilots and this preregistered study was via Prolific. The average cost per completed garden was similar with the two methods, but Prolific recruitment was finally selected in part because paying users results in a more predictable number of gardens completed over shorter spans of time.

The preregistered intention was to update the model with "all UK garden data added [to the system] during the busiest five days of use during May 2025." Busy days were generated via Prolific recruitment. We aimed to collect 14 gardens per working day in the week-long period beginning the 20th of May, and overrecruited by one, resulting in 71 gardens. No useable data not sourced via Prolific recruitment was contributed during this period. All garden data that logically and ethically could be included in model updating was included. There were two reasons why some garden data could not be included, unanticipated in the preregistration: one garden suffered from a technical issue related to measuring distances to buildings in Northern Ireland (see Variables sub-section below), and for five gardens, participants withdrew consent for processing of their data using the Prolific task-return function.

The Prolific internal screening system was used to recruit only UK residents with an upper secondary education, scoring at least 4 on a 1-to-5 environmental concern scale, and with some touch-typing skills. These requirements were intended to optimise motivation and ability to use the system (touch-typing was the Prolific screener most reflective of moderate technical ability) without making the recruitment pool very narrow in comparison to the national population. As further data quality control measures, targeted users were also required to have completed at least five previous Prolific tasks with a minimum of 98% past tasks approved, and to have been born in the UK (systematic analysis of pilot data indicated a need for this latter requirement).

Data reported to Prolific by the 71 included participants indicated 79% had university degrees, 56% were female[1], 10% were students, 69% were in full-time work, 10% were in part-time work, and the median age was 39 (lower quartile 34, upper quartile 49.5). The median task time for included gardens was 22 minutes, and the compensation was £3 per included garden, £3 or £1

[1] Prolific demographic data by default includes sex rather than gender.

(depending on apparent level of effort) for good-faith attempts that were insufficient to support model updating, and a smaller variable quantity (depending on time spent) for participants screened-out for bird misidentification. Table S3 summarises the progress of participants through the recruitment and data provision process.

Table S3: Participant data inclusion determination

| Participant category | Notes | Count | % of those fully accepting task |
|---|---|---|---|
| Initially accepted task | Accepted the task in Prolific, booking a place but not representing a commitment to complete. | 169 | 122% |
| Fully accepted task | Completed the consent and screening questionnaire. Withdrawal here likely prompted by the additional task information given, or poor confidence in identifying birds. | 139 | 100% |
| Screened out for misidentification | Two bird identifications required to be correct. | 29 | 21% |
| Provided insufficient data to support model updating | Despite visiting the app. Many in this category also withdrew. Subcategories below. | 28 | 20% |
| *No data at all* | *(subcategory)* | *20* | *14%* |
| *Drew only a garden boundary* | *(subcategory)* | *4* | *3%* |
| *No sparrow occurrence data* | *(subcategory)* | *4* | *3%* |
| Provided sufficient data but withdrew participation | Data could have been included but consent was withdrawn. Withdrawal here likely reflects concern about garden description quality. | 5 | 4% |
| Described area not a garden | As obviously indicated by aerial photography. | 5 | 4% |
| In Northern Ireland with unmapped buildings | Proper processing impossible for technical reasons (see Variables section below). | 1 | 1% |
| Provided data included in model updating | Sparrow occurrence and at least some habitat data provided. | 71 | 51% |

*Note*: Categories are mutually exclusive except for the subcategories.

The Prolific task description and the screening questionnaire are in the Recruitment Materials section. A data file with demographic data and garden metadata is included in the data package.

###### 4.2.2.2. Non-automatic assistance to participants

Three users reached out via the Prolific messaging system to ask for personal assistance; in each case, attempts were made at assistance, but users changed no data following assistance.

##### *4.2.3. Garden Advice app*

###### 4.2.3.1. Calculation of sparrow scores

For reasons including low sparrow prevalence at cell level, and the simple linearity of the model regarding proximity variables, raw model predictions are not suitable to be presented to users as sparrow scores without transformation. A binary logistic model will tend to output a mean probability modelling the prevalence at data-point level. This mean probability is not ideal for direct presentation to system users as a sparrow score, because the prevalence at data-point level (2 $m^2$ grid cell) is low, although it is high at location level (main text Table 2). For EDS-related model prediction in Garden Advice, we therefore adjust the model intercept to output a mean probability of 50%, so we can explain to system users that "Sparrow scores" of 50% represent averagely good sparrow habitat. For EDS model predictions, we also capped distance variables at values beyond which the GHSP data could not offer reliable predictions, for lack of data (exactly how is described in comments and code in the R script in the preregistration). Otherwise, given beneficial proximity effects and the simple linear nature of the model, cells far from hedges or roofs would always receive a zero score.

The R script that generates model predictions for Garden Advice users operates using a streamlined approach, rather than through the full process of obtaining predictions from an MCMC model object. Because predictions do not use the random model components or express uncertainty, a simple calculation suffices: the sum of the intercept and each predictor variable value multiplied by the relevant fixed coefficient. This approach minimises the latency experienced by the user when requesting model scores – but it does imply that a slower approach would be needed for the expression of uncertainty, which would be beneficial in many contexts.

##### 4.2.3.2. User interface

###### *4.2.3.2.1. System development and basic properties*

The user interface for Garden Advice is characterised by two primary functionalities: (1) drawing polygons on a base map (Google Maps satellite view); (2) obtaining scores representing predictions from models that process the entered map data. Users can enter polygons representing habitats or areas where different animal species are typically observed. Additional functionality includes the ability to enter planning mode. Planning mode copies the existing map to a new version designed to allow what-if edits permitting model score comparisons between the real map and the modified plan.

Garden Advice interfaces were developed over a period of piloting to ensure relative ease of system use. Functionality intended to facilitate the most challenging task of drawing polygons includes multiple available drawing methods (free-hand drawing with mouse or finger or polygon vertex control via tap, click, or GPS), automatic polygon tidying functions, and tutorials. The app also features a Task Panel system enabling users to be guided through a specific sequence of tasks necessary for a particular purpose. The easiest way to understand the Garden Advice interface is to access it at https://garden.earthadvice.org (those providing non-genuine data for testing purposes are asked to enable testing mode). It is also possible to explore the code base – see the “Garden Advice App (REDS_ONE)” sections below.

*4.2.3.2.2. Presentation of sparrow scores*

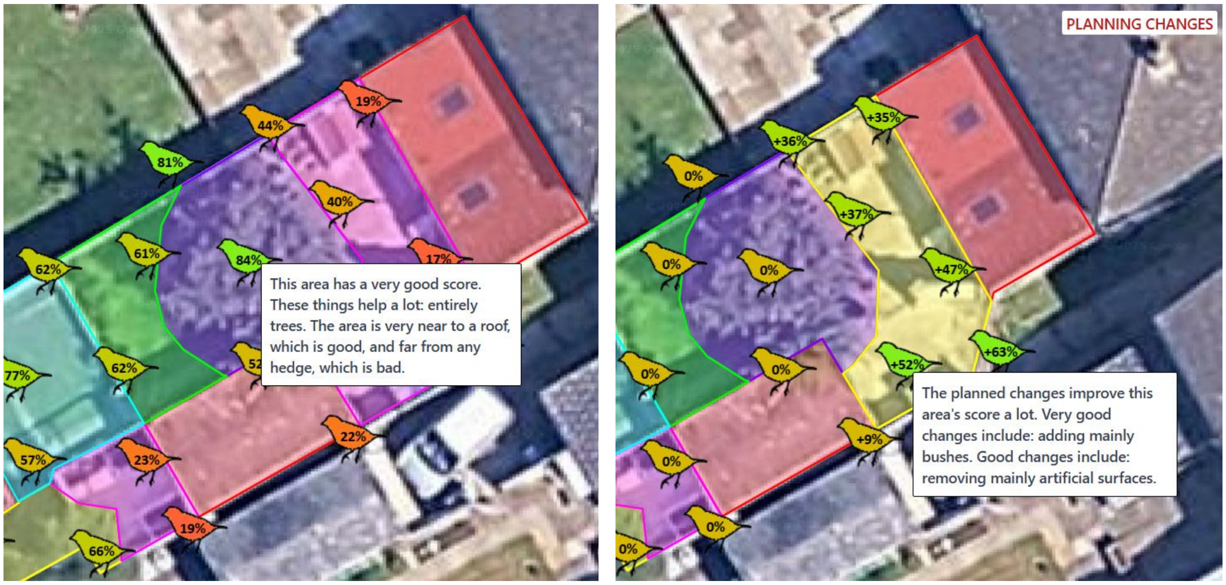

Figure S1. Sparrow scores in standard mode (left) and planning mode (right). The text boxes show explanator texts, generated by routines that turn the regression equation calculations into natural language, available by touching or clicking sparrow score icons. See the main text Figure 3 legend for map colour codes. The base map layer is Google Maps satellite view, © (2025) Google.

When the user chooses to view sparrow scores, they see sparrow icons covering all mapped areas of their garden indicating sparrow scores in percentages (see above for calculation details). Figure S1 (left panel) shows an example garden with sparrow scores. The icon display is zoom sensitive. When zoomed far in, one icon is shown for each 2m grid-cell; when zoomed further out, the icons aggregate to cover broader areas. Figure S1 (right panel) shows an example of what-if analysis using planning mode. In this mode, sparrow score changes caused by planned changes are shown instead of absolute sparrow scores. In this case, simulating replanting of an area of artificial surface with bushes improves the sparrow score.

##### 4.2.3.3. The carbon model

See the Garden Advice Carbon Model top-level section.

#### *4.2.4. The sparrow habitat association model*

##### 4.2.4.1. Variables

*4.2.4.1.1. Distance calculations*

All GHSP cells contained values for both distance variables, with the means and SDs for both almost exactly 0 and 1 respectively, indicating they had been z-transformed (deviations from exactly 0 and 1 are presumed accountable for by subsampling following transformation). To reconstruct approximate distances in metres, we utilised median garden sizes in the postcode areas named in the GHSP data, available from the UK Office for National Statistics (ONS, 2020), and simple assumptions about how buildings and hedges tend to be located in gardens in Glasgow suburbs. Full details are in comments and code in the R script in the preregistration.

Exact GA building distance data was missing in only one case, although users frequently omitted buildings from their maps, because GA imports building location from a publicly available map at individual building scale (Ordnance Survey, 2025). This map is available everywhere in the UK except Northern Ireland, so the one exception was a Northern Ireland garden where the user had not mapped buildings, which was necessarily excluded because building distance could not be set in a way compatible with other GA data. For gardens with no mapped hedges, a different approach was taken, because no external hedge map was available: the distance to the nearest hedge was set to the maximum distance recorded at time of model construction in any GA map with a hedge.

##### 4.2.4.2. Model specification and updating

###### *4.2.4.2.1. Offsetting for case-control data*

Case-control data can be made compatible with non-case-control data in binary logistic regression by offsetting (Huang & Pepe, 2010). All data can be included in one model when an offset is applied to the linear predictor for the case-control data only, which is logit($p$) = ln[$p/(1-p)$], where $p$ is the natural prevalence. For the initial model placed in the GA system (trained only with GHSP data), we therefore estimated natural prevalence from pilot data and offset every data point; in later models including GA data, GHSP but not GA data points were offset. Prevalence is calculated as the grand mean of each location’s cell mean in GA data.

We did not preregister to change this offset as our estimate of prevalence improved with additional GA data, but we deviated from preregistration by changing this in the final GA-updated model, as this is an obvious choice. We also deviate from preregistration by comparing the final GA-updated model with a recreation of the original GHSP-only model that uses an offset calculated from final prevalence data. The effectively gives the GHSP-only model an unrealistic advantage, because the GHSP-only model then incorporates the prevalence data present only in the GA data. We chose this because otherwise any model calibration improvement following updating might reflect nothing more than the trivial fact that the GHSP-only model is uncalibrated for prevalence due to the GHSP case-control data sampling method.

###### *4.2.4.2.2. Model run-time – the slowness of MCMC with large data quantities*

Preregistration stated that models would be fitted using the standard Markov chain Monte Carlo (MCMC) approach with the R package rstanarm (Goodrich et al., 2025). However, we had underestimated how run time for MCMC Bayesian updating would scale up with additional data. It became evident that leave-one-group-out cross-validation (requiring one model to be constructed per garden or colony) was only practical with a faster alternative fitting method – we used Integrated nested Laplace approximation (INLA) via the R-INLA package (Rue et al., 2017). Other analyses are presented using both approaches.

Whereas the day one update with 46 locations took 20 minutes, the day five update with all 103 locations took 8 hours, implying leave-one-out cross-validation would take 34 days. Further, the quadratic scaling implied by the run-time data (Figure S2) indicates that collecting another batch of GA data of the same size and updating the model would take 1.6 days just for the update, with no cross-validation. We did not anticipate this issue at the point of design but now suggest that a REDS implementation using MCMC is impractical, making INLA very important. Future directions for technical implementation may also include methods that do in fact allow model updating without reprocessing all data, such as particle filter approaches (Godsill, 2019).

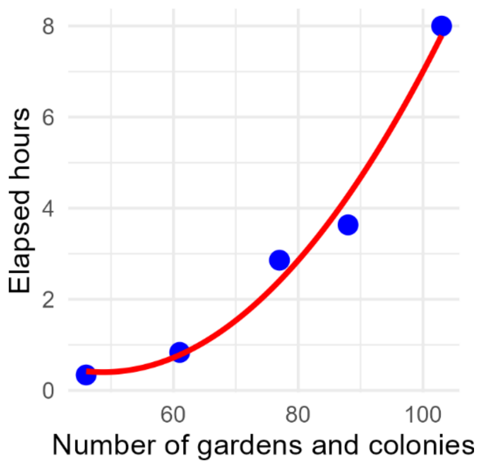

Figure S2. Run time for each of the five model-updating processes during the study period, with a quadratic model fit. The machine running the updating was a consumer Windows laptop workstation purchased in 2025.

*4.2.4.2.3. Prior specification for random slope variances*

Initial attempts to fit INLA models to the final dataset showed poor convergence with MCMC models. However, after model diagnostics suggested issues with the random slopes, we investigated INLA random slope variance priors that were less vague than default and specified comparatively high variance, resulting in good convergence of the INLA models with the MCMC models.

In INLA, random effect variance components are parameterized through their precision ($\tau = 1/\sigma^2$), where σ is the standard deviation. The default prior in R-INLA for random effect precision is Gamma(1, $5\times10^{-5}$) (Gómez-Rubio, 2020), which is very vague, placing minimal constraints on the precision parameter. Initial INLA model fitting attempts using default priors resulted in fixed posteriors that were quite different to the more trustworthy MCMC estimates. We therefore replaced the default prior with penalized complexity (PC) priors (Fuglstad et al., 2019) for random slope variance components. The PC prior is specified through two parameters (u, α) that encode the statement $P(\sigma > u) = \alpha$, where σ is the standard deviation of the random effect. We set $u = 1$ and $\alpha = 0.5$, corresponding to the prior belief that there is a 50% probability that the random slope standard deviation exceeds 1. This represents a prior belief in a relatively high probability of a large amount of variation, which is defensible.

In retrospect, a prior belief in greater variance would have been more appropriate than the defaults (Gelman, 2006), even given no data: random slopes in our model capture between-location variation in habitat associations, which we would expect to be substantial given that (a) inter-location variance in how sparrows use habitats is likely, and (b) spatial autocorrelation within locations would strongly amplify these location-specific patterns.

*4.2.4.2.4. Model updating procedure*

An implication of Bayes theorem is that creating a model with dataset *A* (and a given prior) and then updating it with dataset *B* is equivalent to creating a model with datasets *A* and *B* combined (with the given prior). A practical implication of this is that implementations of Bayesian models frequently don't in practice allow updating of existing model objects. Model updating was therefore always conducted by recreating the model with all data and the original prior.

Garden Advice model updating was conducted through a fully automated once-a-day process where components of the system coordinated to (1) identify new data since last update, (2) convert the vector format to the cell format for new data, (3) deliver this data to a model updating process which runs a model-updating R script to produce a new model-prediction R script containing the new model coefficient values, and (4) install this new prediction script back into the score generating system component. For convenience of monitoring, the model updating process itself was run on a local workstation which automatically communicated with the other components in the cloud via an API (see top-level section "Garden Advice App (REDS_ONE) Technical Architecture Description").

##### 4.2.4.3. Model predictions and their evaluation

###### *4.2.4.3.1. Brier score calculations*

We deviated from the preregistered method for calculating Brier scores (use of the R function scoring::brierscore(), Merkle & Hartman, 2018) for two reasons. Firstly, data from one garden reliably caused that function to throw a rare error for reasons that could not be identified. Secondly, Brier score decomposition requires predicted probabilities to be grouped into bins, with bin boundaries affecting results (Stephenson et al., 2008), but that function allows limited control over the binning approach, with no adaptation to the obtained range of predicted probabilities. Our implementation uses 10 equal-sized bins, adapted individually for each garden, ranging from the minimum to maximum obtained predicted probabilities for that garden. This decision was prompted as follows. First, triage of the error reliably thrown by scoring::brierscore() indicated it would be difficult to debug, so new code was written to implement Brier scores and their decomposition. This led to a focus on the issue of binning, and a realisation that the only approach provided by scoring::brierscore(), rounding of probabilities to the nearest multiple of a given decimal, was not optimal for our gardens, because the ranges of predicted probabilities were highly variable. Our approach of dynamic bin sizes (with the number of bins, 10, chosen to match the mean data points per garden, 469) was the first approach we adopted, and we did not explore further approaches.

Note that gardens that have no variation in sparrow presence (present or absent in every cell) have no defined Brier score decomposition but do have Brier scores.

###### *4.2.4.3.2. Cross-validation*

We used a leave-one-group-out cross-validation process (Vehtari, 2025) which means that a model was never tested for predicting data with which it was trained. This is appropriate for rigorous model validation, because it removes any risk of over-fitting. The process was as follows. For every location (garden or colony) in the prediction dataset, a version of the model was constructed omitting that location's data in training. The predictions for that location were therefore from a model that includes all other training data, but not that location.

###### *4.2.4.3.3. Construction of credible intervals for Brier score change at updating*

Brier scores exist on a per-data-point basis. A difference score was calculated for each data point, as the difference between the predictions from the initial and updated model versions. A Bayesian model was constructed (with stan_glmer() and default parameters) modelling this difference score as nothing but a fixed intercept and random intercepts for each location. The posterior distribution of the intercept shows the strength of the evidence for model differences. Brier decompositions exist on a per-location basis. They were therefore treated in the same way except with a non-random intercept-only stan_glm() model with one difference score per location (probably explaining the reduced power for decompositions suggested by Figure 5).

*4.2.4.3.4. Calculation of effect consistency*

Effect consistency represents the proportion of locations (gardens and colonies) expected to show effects in the direction predicted by the population-level (fixed) effect. This metric accounts for effect heterogeneity, i.e., variation in effect magnitude and direction across locations.

The model assumes location-specific effects follow a normal distribution with mean equal to the fixed effect ($\beta$) and standard deviation equal to the between-location random slope SD ($\sigma$). Under this assumption, the proportion of locations with effects in the predicted direction is $\Phi(\beta/\sigma)$, where $\Phi$ is the cumulative distribution function of the standard normal distribution.

When many gardens lack within-garden variance for a predictor, the hierarchical model has limited information to estimate between-garden variability in random slope, leading to appreciable risk of underestimation of random slope SDs. This was a particular problem for hedge proximity, where only 45% of gardens were informative, and the hyperparameter SD (median 0.501) therefore underestimated the empirical SD calculated in the normal manner from informative gardens (median 0.814). For all effect consistency estimates, we therefore used empirical SDs calculated directly from posterior draws of garden-specific slopes for informative gardens only (within-garden variance > 0).

#### 4.3. Results

##### *4.3.1. Data description*

The median size of holdings mapped in Garden Advice (GA) was 509 $m^2$, first to third quartiles 199 $m^2$ to 1257 $m^2$, mean 1299 $m^2$ (including mapped buildings). Descriptive statistics for model variables are in Table S4.

Table S4. Descriptive statistics for model variables. GA and GHSP are most appropriately compared using cells with no sparrows, given the different sampling approaches.

| | All map cells | | | Map cells with no sparrows | | | | | |
|---|---|---|---|---|---|---|---|---|---|
| | GA | | | GA | | | GHSP | | |
| Variable | Mean | Median | Present | Mean | Median | Present | Mean | Median | Present |
| | | | | Cell percentage cover | | | | | |
| Sparrow | 22% | 17% | 85% | - | - | - | - | - | - |
| Roof | 12% | 4% | 56% | 13% | 4% | 56% | 22% | 20% | 100% |
| Grass | 28% | 26% | 83% | 25% | 24% | 80% | 25% | 23% | 97% |
| Bush | 17% | 14% | 82% | 15% | 9% | 77% | 7% | 4% | 78% |
| Tree | 16% | 7% | 61% | 15% | 6% | 58% | 5% | 2% | 50% |
| Hedge | 3% | 0% | 20% | 3% | 0% | 17% | 7% | 6% | 78% |
| Artificial | 23% | 20% | 85% | 26% | 20% | 85% | 33% | 33% | 100% |
| | | | | Distance to nearest (m) | | | | | |
| Roof | 8.9 | 6.2 | 100% | 8.5 | 5.7 | 100% | 11.4 | 11.1 | 100% |
| Hedge | 14.2 | 16.0 | 20% | 15.6 | 16.8 | 20% | 13.4 | 12.8 | 100% |

*Notes*: For cover variables (including sparrow observation areas), present means the percentage of locations (gardens or colonies) with non-zero cover. For distance variables, present means the percentage of locations where the distance variable was available.

##### *4.3.2. Effect of model updating on model parameters*

###### 4.3.2.1. Convergence of INLA with MCMC

Main text Figure 4 indicates that INLA approximations converged very well with MCMC models, with one exception – the hedge proximity parameter. As even this parameter converged reasonably well, INLA appears a very good approximation in this context. The imperfect convergence for hedge proximity likely relates to greater difficulty fitting hedge parameters due to the absence of hedges from most GA gardens.

This is unsurprising given that hedges were reported present in only 20% of GA gardens, whereas all other features were reported present in most gardens (main text Table 3). When hedges were absent, the maximum observed distance was imputed for this variable, which causes a total lack of heterogeneity in this variable within most gardens, and so also low mean heterogeneity between gardens. This would present an appreciable challenge to estimation of the random slope hyperparameters, and thus also the corresponding fixed parameter, with these problems potentially exacerbating one another. This problem can be expected to be more acute for INLA than MCMC as it is an approximate method.

##### *4.3.3. Effect of model updating on predictive power*

###### 4.3.3.1. Spatial autocorrelation testing and leave-one-region-out cross-validation

If the relationship between habitat and sparrow presence, or habitat itself, varies spatially, with nearby locations more similar than distant ones (spatial autocorrelation), then a cross-validation excluding nearby GA gardens would be an appropriate robustness check. We began to examine this issue by constructing a correlogram, which plots Moran’s I (a measure of correlation) at different spatial lags (the correlogram uses residuals from fixed-effect-only predictions for GA gardens from the updated MCMC model). Higher similarity between nearby gardens would be indicated by Moran’s I values that begin greater than zero and decrease with lag. The lack of any such pattern, with Moran’s I consistently trending around zero at all distance lags (Figure S3), indicates a lack of such autocorrelation (Sokal & Oden, 1978).

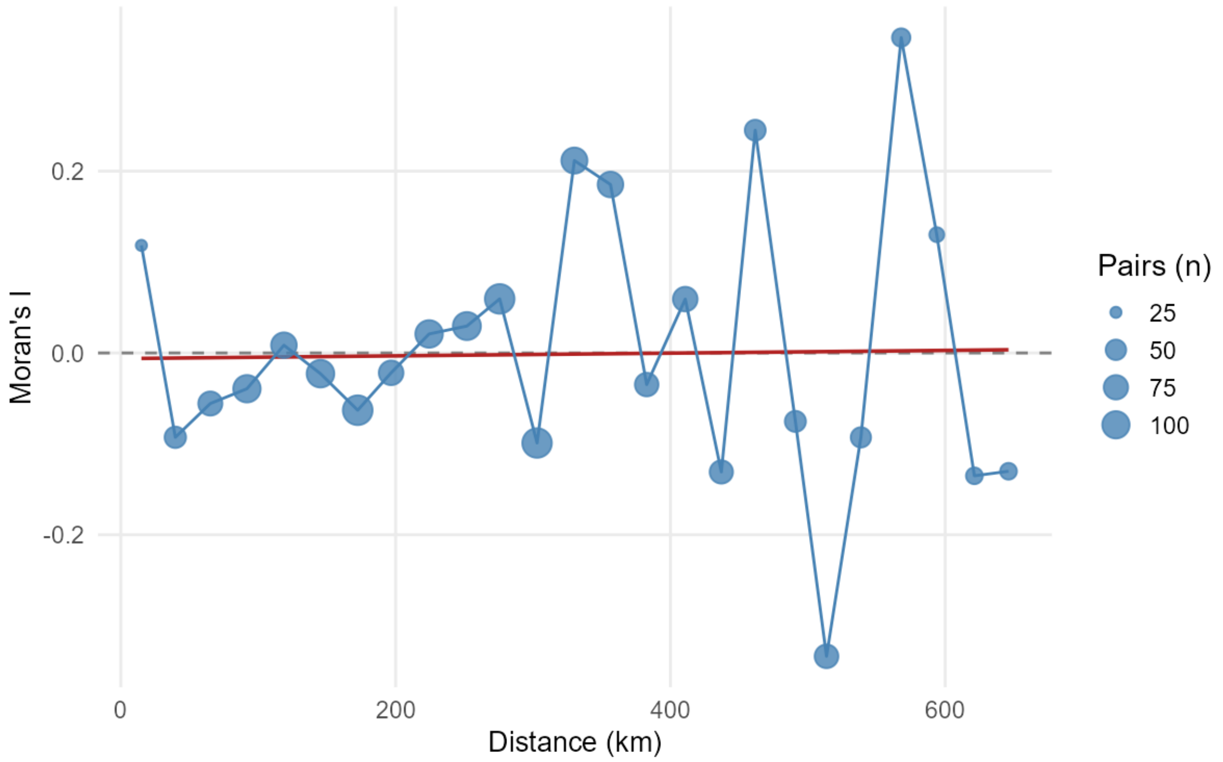

Figure S3. Spatial correlogram of fixed-effect residuals. Point size shows the number of garden pairs per distance bin; the red line shows the linear trend weighted by pairs per bin. The bin width is 27 km (the median nearest neighbour distance). The distance cutoff is 669 km (the 75$^{th}$ percentile pairwise distance).

Nevertheless, an alternative leave-one-out approach would provide a robustness check, and the larger the areas left out, the greater the robustness. We therefore divided the UK into four large regions by combining the smaller standard UK regions (South of England: South West, South East, and London; Wales and the English Midlands: Wales, West Midlands, East Midlands, and the East of England; North of England: North West, North East, and Yorkshire and The Humber; Scotland and Northern Ireland: Scotland and Northern Ireland). The leave-one-out analysis was conducted using the same approach as the preregistered leave-one-garden-out analysis (Section 4.2.4.3), except that four models were constructed (one to leave out each region rather than garden) and predictions for each garden were obtained from the model which left out its region. Mean overall Brier scores are reported in the main text (they do not differ).

##### 4.3.3.2. Additional binary discrimination statistics

To examine model performance when thresholded to produce binary predictions, we first optimised the threshold to match the predicted prevalence to observed prevalence (an approach which is generally recommended in ecology, Freeman & Moisen, 2008). We examine the Matthews correlation coefficient (MCC), a metric superior to other binary discrimination metrics (Chicco & Jurman, 2023), along with a more intuitive metric, the positive likelihood ratio. When optimising threshold for prevalence, using MCMC, the GHSP-only model predicts sparrows poorly in the GA dataset, whereas the GA-updated model shows a modest improvement (Table S5). When using INLA, updating results in a similar improvement (Table S6); when optimising for MCC rather than matching predicted and observed prevalence, the GHSP-only model again shows poor discrimination, but the GA-updated version is markedly improved (Table S7).

Table S5. MCMC model binary discrimination performance, before and after model updating

| | | GHSP-only model | | GA-updated model | |
|---|---|---|---|---|---|
| Threshold optimised for prevalence | | 0.72 | | 0.19 | |
| *Confusion matrices* | | | | | |
| | | Predicted | | Predicted | |
| | | No sparrow | Sparrow | No sparrow | Sparrow |
| Observed | No sparrow | 61.9% | 16.4% | 62.8% | 15.5% |
| | Sparrow | 16.4% | 5.3% | 15.6% | 6.1% |
| *Discrimination metrics* | | | | | |
| False positive rate (FPR) | | 21.0% | | 19.8% | |
| True positive rate (TPR) | | 24.3% | | 28.0% | |
| Positive likelihood ratio (TPR/FPR) | | 1.16 | | 1.42 | |
| Mathews correlation coefficient (MCC) | | 0.03 | | 0.08 | |

*Notes*: Prediction is for GA data. The discrimination threshold is optimised separately for each model, so the performance comparison is valid. The confusion matrix is the mean of the confusion matrices for all GA gardens.

Table S6. INLA model binary discrimination performance, before and after model updating

| | | GHSP-only model | | GA-updated model | |
|---|---|---|---|---|---|
| Threshold optimised for prevalence | | 0.66 | | 0.41 | |
| *Confusion matrices* | | | | | |
| | | Predicted | | Predicted | |
| | | No sparrow | Sparrow | No sparrow | Sparrow |
| Observed | No sparrow | 62.0% | 16.3% | 62.8% | 15.5% |
| | Sparrow | 16.0% | 5.7% | 15.2% | 6.5% |
| *Discrimination metrics* | | | | | |
| False positive rate (FPR) | | 20.9% | | 19.8% | |
| True positive rate (TPR) | | 26.4% | | 29.9% | |
| Positive likelihood ratio (TPR/FPR) | | 1.27 | | 1.51 | |
| Mathews correlation coefficient (MCC) | | 0.06 | | 0.10 | |

*Notes*: Prediction is for GA data. The discrimination threshold is optimised separately for each model, so the performance comparison is valid. The confusion matrix is the mean of the confusion matrices for all GA gardens.

Table S7. MCMC model binary discrimination performance, before and after model updating

| | | GHSP-only model | | GA-updated model | |
|---|---|---|---|---|---|
| Threshold optimised for MCC | | 0.39 | | 0.41 | |
| *Confusion matrices* | | | | | |
| | | Predicted | | Predicted | |
| | | No sparrow | Sparrow | No sparrow | Sparrow |
| Observed | No sparrow | 34.0% | 44.3% | 71.9% | 6.4% |
| | Sparrow | 7.9% | 13.8% | 17.5% | 4.2% |
| *Discrimination metrics* | | | | | |
| False positive rate (FPR) | | 56.5% | | 8.1% | |
| True positive rate (TPR) | | 63.5% | | 19.3% | |
| Positive likelihood ratio (TPR/FPR) | | 1.12 | | 2.37 | |
| Mathews correlation coefficient (MCC) | | 0.06 | | 0.15 | |

*Notes*: Prediction is for GA data. The discrimination threshold is optimised separately for each model, so the performance comparison is valid. The confusion matrix is the mean of the confusion matrices for all GA gardens.

##### *4.3.4. Effect consistency*

###### 4.3.4.1. Effect consistency for the initial GHSP-only model

Figure S4 shows the equivalent of the main text effect consistency plot (Figure 6), but for GHSP-only data rather than after updating with GA data. It clearly indicates that GHSP data by itself is generally insufficient to support confident predictions, except that roof proximity is beneficial and artificial surface is very likely detrimental.

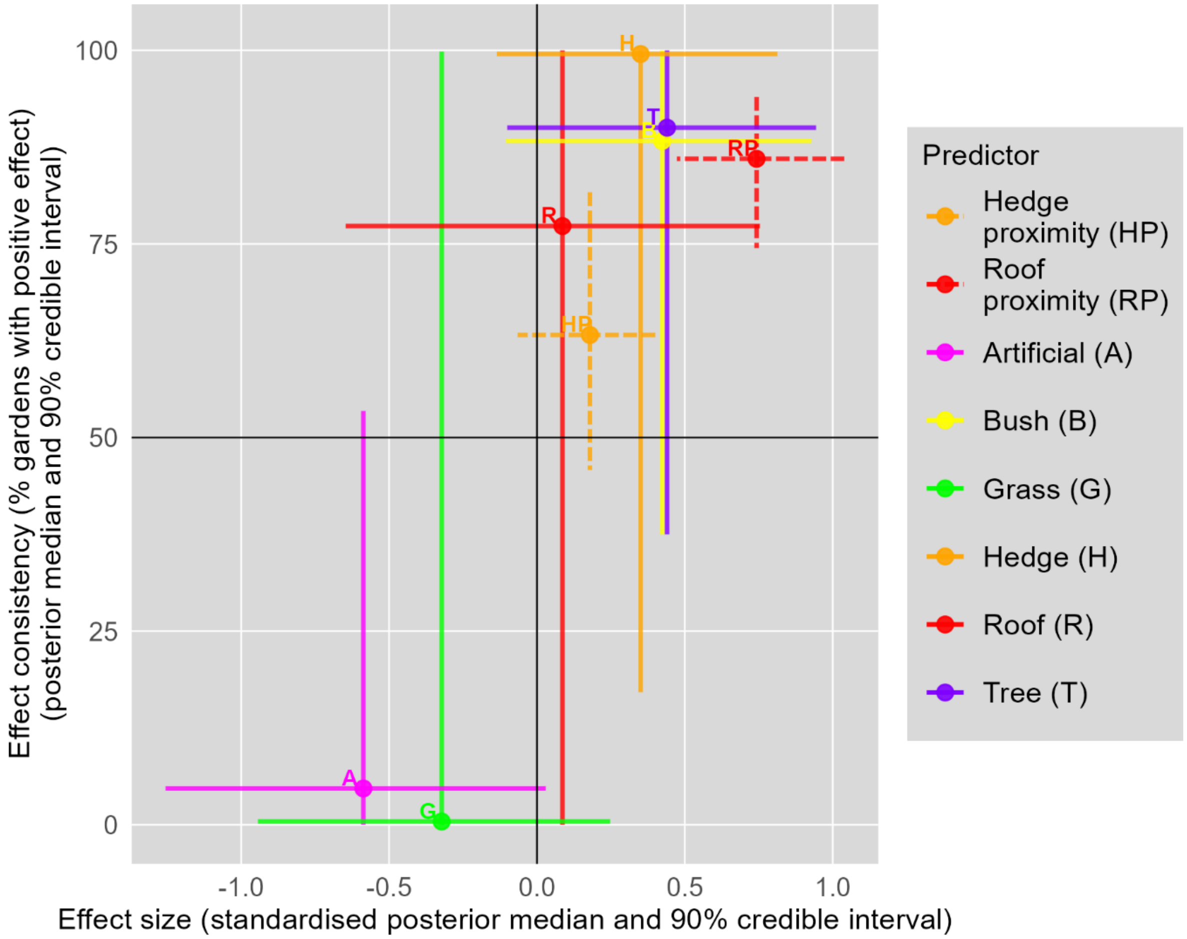

Figure S4. Effect consistency and standardised effect size across predictors for GHSP-only MCMC model. Effect sizes are standardized by multiplying regression parameters by the pooled within-garden (or colony) standard deviation for each predictor.

#### *5. Data and Code Package*

This package is available at the repository https://github.com/artifact-u17x/REDS_ONE_POC1.

It contains the top-level directories:

- subsequentAnalysis: The R and Python scripts which run the analyses reported in the manuscript. All the primary analyses are begun in analysis.R; Python is used only for the UK map figure. Because some models are slow to run (even the cross-validation, although it is INLA, because it creates many models), we include saved model objects which by default are reloaded instead of recreated.
- fullProofOfConcept: The scheduled batch scripts and R scripts which ran the daily model updating during the study model updating period. The process also uses the file commonCode.R in the repository root. Note that the datafile subdirectory cellingBatches is not present in the shared repository because this raw data cannot be shared for reasons of privacy; instead, a processed version is made available in the subsequentAnalysis directory which is automatically loaded by the analysis scripts.
- additionalData: Data files describing participant demographics and in-app behaviour.

- redistributedData: The GHSP data file, redistributed as allowed under its own license (Matthiopoulos et al., 2019).
- searchingForOtherREDS: The database and a processing R script relating to our systematic search for other REDS systems.

Large Language Models (LLM) were used during creation of much of the code in this package, for generating code from scratch and for modifying and debugging (including Anthropic's Claude Sonnet 4.5). The authors have manually checked all LLM generated code and take responsibility for it. We do not separately label code as originally produced by human or LLM, because LLM-produced code is frequently subsequently human-modified, and vice versa, so line-by-line labelling would be required.

### *6. Garden Advice App (REDS_ONE) Technical Architecture Description*

#### 6.1. System Overview

REDS_ONE (Reciprocal Environmental Decision Support One) is the first known REDS implementation supporting decisions about ecosystem management. Garden Advice is a particular instantiation of REDS_ONE which is itself a generalised application class allowing implementation of different REDS applications. REDS_ONE produces enterprise web applications enabling nature users to create maps of outdoor spaces, collect ecological data, and view analytical model results. The platform provides interactive mapping tools for end users, an administrative dashboard for scientists to view data and manage models, and flexible configuration for different mapping scenarios. Production and staging environments support live and testing operations.

#### 6.2. Infrastructure & Hosting

**AWS Cloud Computing** provides the complete hosting infrastructure with reliability, scalability, and integrated backup solutions.

**Container orchestration: Docker Compose** orchestrates containerised services including web applications, databases, and model execution environments, ensuring consistent deployment across all environments.

**Compute:** The production environment has two T4G Small **EC2** instances (ARM-based Graviton processors) behind an AWS Elastic Load Balancer (**ELB**), providing horizontal scaling capability and high availability. AWS Security Groups deliver firewall protection and network isolation.

**Web Server: NGINX** handles high-performance static file serving, SSL/TLS termination, request routing, and efficient connection handling.

#### 6.3. Application Architecture

##### *6.3.1. Backend: Laravel (PHP)*

**Laravel**, built and programmed in **PHP**, provides the business logic layer with:

- **Asynchronous Task Handling:** Queue system manages long-running model executions without blocking user requests

- **Authentication & Authorization:** Built-in user management for administrators and scientists
- **API Layer:** RESTful endpoints for client communication
- **Data Management:** Tools for viewing/downloading user data and model results
- **Model Administration:** Interface for managing and deploying analytical models

##### *6.3.2. Frontend: Svelte (JavaScript)*

**Svelte** lets developers write **JavaScript** (along with **HTML** and **CSS**), which its compiler transforms into highly optimised JavaScript for the user-facing application. It powers the user-facing application with:

- **Fat Client Design:** Rich interactivity with minimal server round-trips
- **API-Driven:** REST API communication with Laravel backend
- **OpenLayers Integration:** Professional mapping library for spatial features
- **Generalised Design:** Easily configurable for different scenarios

Advantages include compile-time optimization, reactive data binding, smaller bundle sizes, and excellent performance for complex mapping interactions.

#### 6.4. Data Management

**PostgreSQL with PostGIS** hosted on **AWS RDS** provides:

- **Spatial Data Support:** Storage and querying of geographic features
- **Relational Integrity:** ACID compliance ensures data consistency
- **AWS RDS Benefits:** Automated backups, point-in-time recovery, managed updates, high availability, performance monitoring

Stores user-created maps (spatial geometries), ecological observations, model inputs/outputs, and user accounts/permissions.

#### 6.5. Analytical Models

**Containerised R Runtime** executes models in isolated environments (architecture supports Python, Julia, etc.). **AWS S3 Buckets** provide centralised storage for model scripts with version control and automatic synchronization across instances.

**Queue-Based Execution:** Laravel queue system manages model run requests asynchronously, preventing UI blocking while supporting progress tracking, result notification, and concurrent model runs.

**Model Management:** Scientists install new models through the admin dashboard or automatically via API with automatic deployment to execution containers, version tracking, and rollback capability.

##### 6.6. Development & Deployment

**Git with GitHub** manages version control, collaboration, branch-based development, code review, and issue tracking. Automated deployment via git push with post-receive hooks for minimal-downtime deployments, consistent process, rapid iteration and easier rollback.

##### 6.7. Key Technical Strengths

**Scalability:** Horizontal scaling via load-balanced EC2 instances, containerised services, queue-based model execution

**Reliability:** High availability through redundant instances, automated database backups, load balancer health checks

**Flexibility:** Generalised frontend architecture, multi-language model runtime support, configuration-driven deployment

**Performance:** Asynchronous model execution, efficient client-side rendering, optimised spatial queries, static asset caching

**Maintainability:** Clear separation of concerns, containerised environments, automated deployment, source control

#### *7. Garden Advice App (REDS_ONE) Code Package*

The REDS_ONE code base, in the form it took at the time of the current study, is shared undocumented and unsupported and for the purpose of academic transparency only, at the repository https://github.com/artifact-u17x/REDS_ONE.

LLM was used during creation of some of the code in this package, for generating code from scratch and for modifying and debugging (including Anthropic's Claude Sonnet 4.5 and multiple versions of ChatGPT). The authors have manually checked all LLM generated code and take responsibility for it. We do not separately label code as originally produced by human or LLM, because LLM-produced code is frequently subsequently human-modified, and vice versa, so line-by-line labelling would be required.

#### *8. Garden Advice Carbon Model*

The model allocates a carbon score to each mapped garden patch (rather than every grid cell, as for the sparrow model). Scores are displayed in hours of human life, using human face icons (Figure S5), and as for sparrow scores, explanator texts are available by tapping or clicking icons.

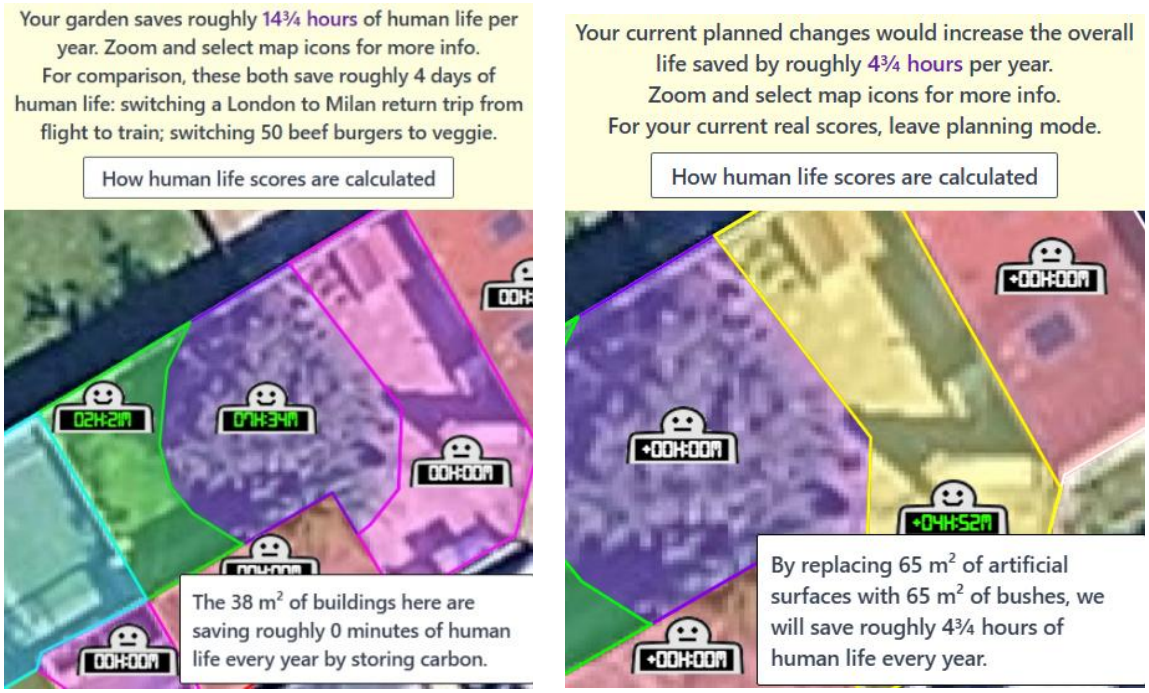

Figure S5. Carbon scores in standard mode (left) and planning mode (right). The text boxes show explanator texts available by touching or clicking score icons. See the main text Figure 3 legend for map colour codes. The base map layer is Google Maps satellite view, © (2025) Google.

To further explain the carbon model, we provide the same explanatory text that is available within the application. We note that we are now in the process of refining our figures for grass carbon sequestration thanks to additional recent review work (Öberg, 2025). The current explanatory text that system users read is as follows (with references added in appropriate format):

The rough amount of human life your garden saves by safely storing carbon pollution is based on two sets of scientific studies.

Corey Flude and colleagues from the University of Guelph reviewed how much carbon is stored per year by different garden plants (Flude et al., 2022). Our specialist examined their findings (see their Table 9) and we have directly adopted most of their figures. They found very similar average values for trees and shrubs, and their results were more reliable for trees due to there being more data, so for hedges and bushes we use the same figure as for trees.

The second set of studies was reviewed in by Joshua Pearce and Richard Parncutt from universities in Canada and Austria (Pearce & Parncutt, 2023). They estimate, with a very wide margin of error, that releasing 1000 tons of carbon leads to one human death this century, through causes such as heat waves, famine due to crop failure, forest fires, floods, and hospital power outages. The most important study (by Daniel Bressler of Columbia University) shows how extreme heat directly kills predictable numbers (Bressler, 2021).

We combine the plant carbon-storage figures with the 1000 ton rule and the pre-pandemic measurement of a human life lasting on average 73 years (World Health Organization, 2024), to find that growing grass saves on average 4 minutes of human life per square metre per year, and trees, bushes, and hedges 7 minutes. Figures can vary quite a lot, and the scores we provide will not apply in all settings.

As the burgers (Saget et al., 2021) and Milan (UIC, 2025) trip comparisons indicate (above), unless you have a very large garden, you can have much more personal impact in other ways. Further, as another study shows (Warburg et al., 2021), thinking about offsetting personal carbon isn't a helpful way to move towards the important changes society needs to make. But, if you do carry out the kind of mini-rewilding project that would improve your garden's carbon life score, it would likely have further benefits, such as helping other local wildlife (sparrows and more).

### *9. Recruitment Materials*

#### 9.1. Study advertisement used within Prolific

We are testing a simple app (no install required) where users describe a garden and find out how it supports life. The goals are scientific research, and information provision to the public.

You will map areas of a residential garden you are very familiar with and have known for at least a year, by marking on a satellite-style map, and report on birds you may have seen there. If the idea of accurately marking a map sounds challenging, this task might not be right for you, as we do require the mapping to be of a minimum standard.

If you are not very familiar with a residential garden in the UK or cannot recognise very common UK garden birds such as blackbird, robin, and house sparrow, you will be screened out with a minimum rate of pay.

If the only garden you are very familiar with is larger than about 500 m2 (roughly two tennis courts) the work may take longer than 15 minutes and so you are not recommended to participate.

Payment will be processed within 24 hours.

We are [REDACTED FOR ANONYMITY].

#### 9.2. Screening questionnaire

**Map a garden you are very familiar with, including visiting birds**

The data we collect in this task will be used as described in the Privacy statement for the app you will use, except that:

- we will not collect any personal information, and because you are anonymous, the data protection (GDPR) sections are not relevant;
- when the app uses the phrase "your garden" it means a garden you are very familiar with, which may or may not be your personal garden, but it must be a residential garden (not a park);
- you must accept the default privacy level "Share with scientists".

Remember that you have committed to accurately mapping your garden. In our experience this is not difficult for most recruited Prolific workers, because the app explains exactly what to do, but if you are concerned, please return the task. Below is an example of what you will create. Note that:

- The whole of the residential garden needs to be covered with shapes (patches) telling us what is there (e.g., bush, grass, tree, building, artificial surface).
- The edges of the patches are in the right places on the aerial photograph.

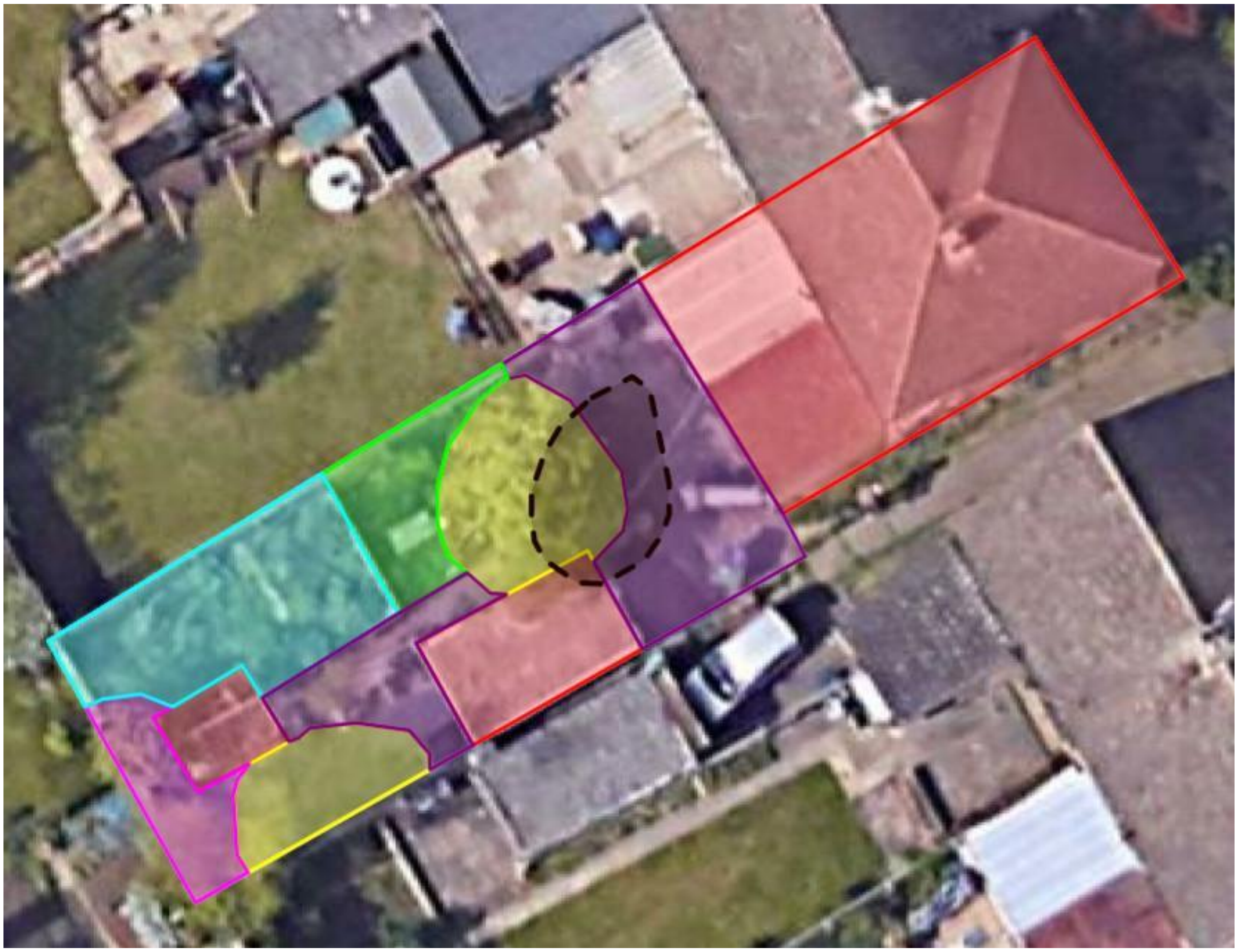

[The satellite view in the above image is © (2025) Google]

In summary: you will accurately map a residential garden you are familiar with, and report birds in it, and we will keep the anonymous data and use it for scientific research which may include sharing it with other scientists.

We are [REDACTED FOR ANONYMITY].

◯ I understand and consent.

◯ I do not consent and so I return the task.

First, we have three questions to confirm you are suitable for this task.

Will you accurately map a residential garden in the UK that you are very familiar with because you have had regular access to it over the whole of the past year?

○ Yes

○ No

Which of these is a blackbird?

○ 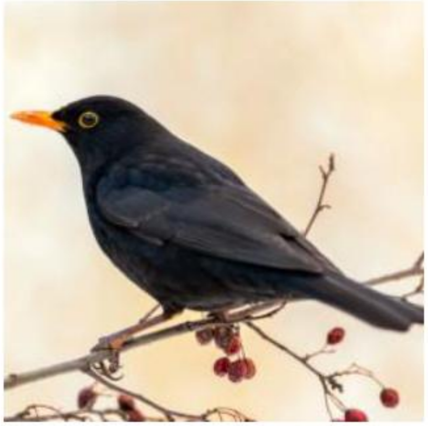

○ 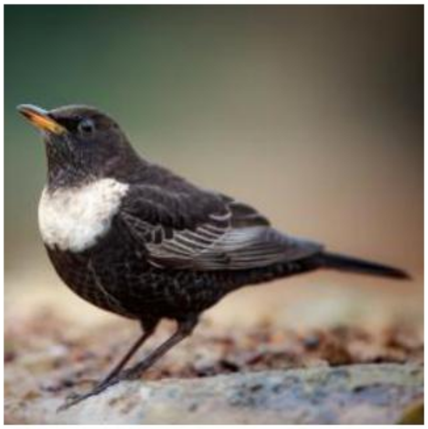

○ 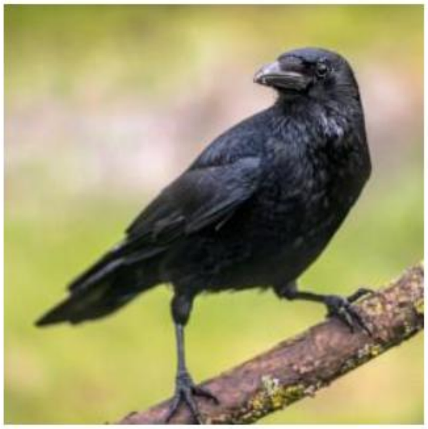

○ 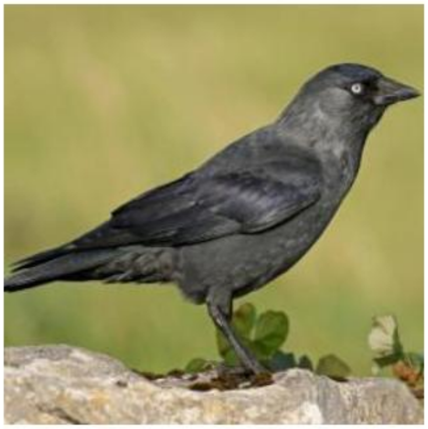

Which of these is a house sparrow?

○ 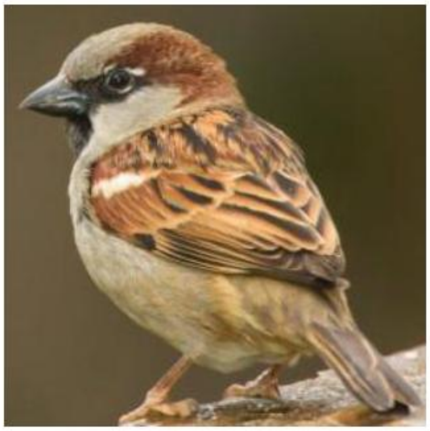

○ 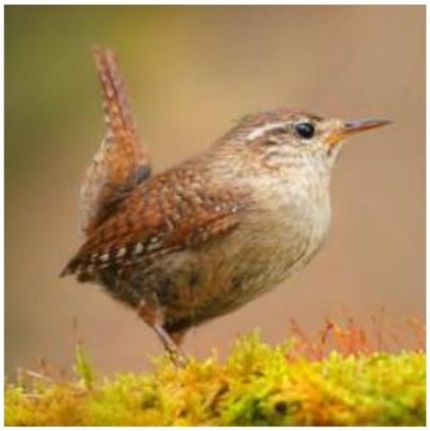

○ 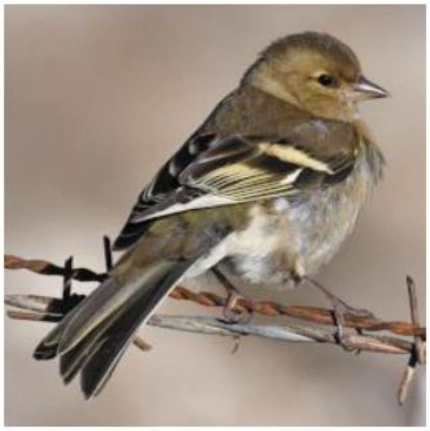

○ 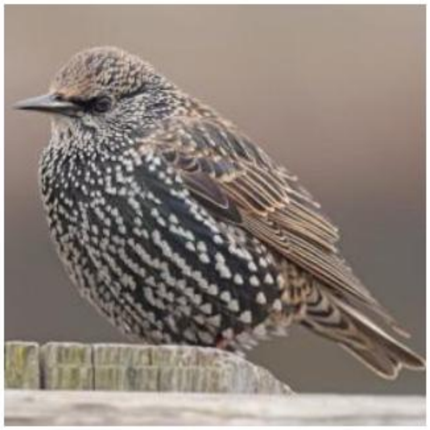

Thank you - your bird identifications were correct and you are suitable for the mapping task. Before the mapping task begins, please answer the following four questions:

Have you ever engaged in nature conservation activities?

○ Yes, on a professional level

○ Yes, on a voluntary basis only

○ No

Have you ever engaged in farming or forestry?

○ Yes

○ No

Have you ever foraged or hunted for wild food, for example fishing to eat or mushroom picking?

○ Yes

○ No

Have you ever grown plants for food?

○ Yes

○ No

You will very shortly proceed to the app.

When you are in the app, pay close attention to the app's task panel (it says "Current Task" and is visible after the intro sequence). You will need to complete every task in the task panel. It's possible to do things in the app that you don't need to do as part of these tasks. It's easiest if you just concentrate on doing the tasks in the task panel, in the order you are asked to do them.

When the final Task is complete, the task panel will show you a completion symbol so that you know you have finished.

Now visit the app by clicking here. Do not use any other link to visit the app. When you see a completion symbol in the app, select it below, and submit this form to return to Prolific where you will be registered as having completed.

○ ★

○ ◆

○ ▲

○ ●

feedback When you have completed the tasks in the app, if you have any time left, please describe here any difficulties you might have had using the app.

________________________________________________________________

________________________________________________________________

#### *10. References*

Bressler, R. D. (2021). The mortality cost of carbon. *Nature Communications*, *12*(1), 4467. https://doi.org/10.1038/s41467-021-24487-w

Chicco, D., & Jurman, G. (2023). The Matthews correlation coefficient (MCC) should replace the ROC AUC as the standard metric for assessing binary classification. *BioData Mining*, *16*(1), 4. https://doi.org/10.1186/s13040-023-00322-4

Cornell Lab of Ornithology. (2025). *Identify bird songs and calls with Sound ID. Cornell Lab Merlin Bird ID*. Retrieved 2025-12-14 from https://merlin.allaboutbirds.org/sound-id/

De Koning, K. (2025). The crane radar: Development and deployment of an operational eco-digital twin. *Ecological Informatics*, *85*, 102938. https://doi.org/https://doi.org/10.1016/j.ecoinf.2024.102938

Flude, C., Ficht, A., Sandoval, F., & Lyons, E. (2022). Development of an Urban Turfgrass and Tree Carbon Calculator for Northern Temperate Climates. *Sustainability*, *14*(19), 12423. https://www.mdpi.com/2071-1050/14/19/12423

Freeman, E. A., & Moisen, G. G. (2008). A comparison of the performance of threshold criteria for binary classification in terms of predicted prevalence and kappa. *Ecological Modelling*, *217*(1), 48-58. https://doi.org/https://doi.org/10.1016/j.ecolmodel.2008.05.015

Fuglstad, G.-A., Simpson, D., Lindgren, F., & Rue, H. (2019). Constructing Priors that Penalize the Complexity of Gaussian Random Fields. *Journal of the American Statistical Association*, *114*(525), 445-452. https://doi.org/10.1080/01621459.2017.1415907

Gelman, A. (2006). Prior distributions for variance parameters in hierarchical models (comment on article by Browne and Draper). *Bayesian Analysis*, *1*(3), 515-534, 520. https://doi.org/10.1214/06-BA117A

Godsill, S. (2019, 12-17 May 2019). Particle Filtering: the First 25 Years and beyond. ICASSP 2019 - 2019 IEEE International Conference on Acoustics, Speech and Signal Processing (ICASSP),

Gómez-Rubio, V. (2020). Priors in R-INLA. In V. Gómez-Rubio (Ed.), *Bayesian Inference with INLA*. Chapman & Hall/CRC Press. https://becarioprecario.bitbucket.io/inla-gitbook/index.html

Goodrich, B., Gabry, J., Ali, I., & Brilleman, S. (2025). *rstanarm: Bayesian applied regression modeling via Stan. R package version 2.32.1*. https://mc-stan.org/rstanarm/

Gorleri, F. C., Jordan, E. A., Roesler, I., Monteleone, D., & Areta, J. I. (2023). Using photographic records to quantify accuracy of bird identifications in citizen science data. *Ibis*, *165*(2), 458-471. https://doi.org/https://doi.org/10.1111/ibi.13137

Huang, Y., & Pepe, M. S. (2010). Assessing risk prediction models in case–control studies using semiparametric and nonparametric methods. *Statistics in Medicine*, *29*(13), 1391-1410. https://doi.org/https://doi.org/10.1002/sim.3876

Lefort, T., Affouard, A., Bonnet, P., Charlier, B., Joly, A., & Salmon, J. (2024). Cooperative learning of Pl@ ntNet's Artificial Intelligence algorithm using label aggregation. JDS 2024-55. Journées de Statistique,

Mason, B. M., Mesaglio, T., Barratt Heitmann, J., Chandler, M., Chowdhury, S., Gorta, S. B. Z., Grattarola, F., Groom, Q., Hitchcock, C., Hoskins, L., Lowe, S. K., Marquis, M., Pernat, N., Shirey, V., Baasanmunkh, S., & Callaghan, C. T. (2025). iNaturalist accelerates biodiversity research. *BioScience*, *75*(11), 953–965. https://doi.org/10.1093/biosci/biaf104

Matthiopoulos, J., Field, C., & MacLeod, R. (2019). Predicting population change from models based on habitat availability and utilization. *Proceedings of the Royal Society B: Biological Sciences*, *286*(1901), 20182911. https://doi.org/10.1098/rspb.2018.2911

Merkle, E. C., & Hartman, R. (2018). Weighted Brier score decompositions for topically heterogenous forecasting tournaments. *Judgment and Decision Making*, *13*(2), 185-201. https://doi.org/10.1017/S1930297500007099

Naturalis Biodiversity Centre. (2020). *Om Artsorakelet*. Retrieved 2025-12-14 from https://artsdatabanken.no/Pages/299643

Newman, J. P., Maier, H. R., Riddell, G. A., Zecchin, A. C., Daniell, J. E., Schaefer, A. M., van Delden, H., Khazai, B., O'Flaherty, M. J., & Newland, C. P. (2017). Review of literature on decision support systems for natural hazard risk reduction: Current status and future research directions. *Environmental Modelling & Software*, *96*, 378-409. https://doi.org/https://doi.org/10.1016/j.envsoft.2017.06.042

Öberg, E. (2025). *Kolbindning i gräsmattor : hur skötselåtgärder påverkar kolbalansen i gräsmattor och hur denna kan förbättras* [Honours thesis, SLU, Dept. of Urban and Rural Development]. https://stud.epsilon.slu.se/20992/

ONS. (2020). *One in eight British households has no garden*. Retrieved 2025-05-19 from https://www.ons.gov.uk/economy/environmentalaccounts/articles/oneineightbritishhouseholdshasnogarden/2020-05-14

Ordnance Survey. (2025). *OS OpenMap - Local*. Retrieved 2025-05-19 from https://osdatahub.os.uk/data/downloads/open/OpenMapLocal

Parker, T., Fraser, H., & Nakagawa, S. (2019). Making conservation science more reliable with preregistration and registered reports. *Conservation Biology*, *33*(4), 747-750. https://doi.org/https://doi.org/10.1111/cobi.13342

Pearce, J. M., & Parncutt, R. (2023). Quantifying Global Greenhouse Gas Emissions in Human Deaths to Guide Energy Policy. *Energies*, *16*(16). https://doi.org/10.3390/en16166074

Rue, H., Riebler, A., Sørbye, S. H., Illian, J. B., Simpson, D. P., & Lindgren, F. K. (2017). Bayesian Computing with INLA: A Review. *Annual Review of Statistics and Its Application*, *4*(Volume 4, 2017), 395-421. https://doi.org/https://doi.org/10.1146/annurev-statistics-060116-054045

Saget, S., Porto Costa, M., Santos, C. S., Vasconcelos, M., Styles, D., & Williams, M. (2021). Comparative life cycle assessment of plant and beef-based patties, including carbon opportunity costs. *Sustainable Production and Consumption*, *28*, 936-952. https://doi.org/https://doi.org/10.1016/j.spc.2021.07.017

Sokal, R. R., & Oden, N. L. (1978). Spatial autocorrelation in biology: 1. Methodology. *Biological Journal of the Linnean Society*, *10*(2), 199-228. https://doi.org/10.1111/j.1095-8312.1978.tb00013.x

Stephenson, D. B., Coelho, C. A., & Jolliffe, I. T. (2008). Two extra components in the Brier score decomposition. *Weather and Forecasting*, *23*(4), 752-757. https://doi.org/doi.org/10.1175/WAF1034.1

UIC. (2025). *EcoPassenger*. Retrieved 2025-05-01 from https://www.ecopassenger.org/

Vehtari, A. (2025). *Can cross-validation be used for hierarchical / multilevel models?* https://users.aalto.fi/~ave/CV-FAQ.html#8_Can_cross-validation_be_used_for_hierarchical__multilevel_models

Warburg, J., Frommeyer, B., Koch, J., Gerdt, S.-O., & Schewe, G. (2021). Voluntary carbon offsetting and consumer choices for environmentally critical products—An experimental study. *Business Strategy and the Environment*, *30*(7), 3009-3024. https://doi.org/https://doi.org/10.1002/bse.2785

World Health Organization. (2024). *Life expectancy at birth (years)*. Retrieved 2025-05-01 from https://data.who.int/indicators/i/90E2E48
